# A Comprehensive Genome-wide and Phenome-wide Examination of BMI and Obesity in a Northern Nevadan Cohort

**DOI:** 10.1101/671123

**Authors:** Karen A. Schlauch, Robert W. Read, Vincent C. Lombardi, Gai Elhanan, William J Metcalf, Anthony D. Slonim, the 23andMe Research Team, Joseph J. Grzymski

**Affiliations:** Institute for Health Innovation, Renown Health, Reno, NV, USA; Desert Research Institute, Reno, NV, USA; Department of Microbiology and Immunology, University of Nevada, School of Medicine, Reno, NV, USA; Renown Health, Reno, NV, USA; 23andMe, Inc. Mountain View, CA, USA

## Abstract

The aggregation of Election Health Records (EHR) and personalized genetics leads to powerful discoveries relevant to population health. Here we perform genome-wide association studies (GWAS) and accompanying phenome-wide association studies (PheWAS) to validate phenotype-genotype associations of BMI, and to a greater extent, severe Class 2 obesity, using comprehensive diagnostic and clinical data from the EHR database of our cohort. Three GWASs of 500,000 variants on the Illumina platform of 6,645 Healthy Nevada participants identified several published and novel variants that affect BMI and obesity. Each GWAS was followed with two independent PheWASs to examine associations between extensive phenotypes (incidence of diagnoses, condition, or disease), significant SNPs, BMI, and incidence of extreme obesity. The first GWAS excludes DM2-diagnosed individuals and focuses on associations with BMI exclusively. The second GWAS examines the interplay between Type 2 Diabetes (DM2) and BMI. The intersection of significant variants of these two studies is surprising. The third complementary case-control GWAS, with cases defined as extremely obese (Class 2 or 3 obesity), identifies strong associations with extreme obesity, including established variants in the *FTO* and *NEGR1* genes, as well as loci not yet linked to obesity. The PheWASs validate published associations between BMI and extreme obesity and incidence of specific diagnoses and conditions, yet also highlight novel links. This study emphasizes the importance of our extensive longitudinal EHR database to validate known associations and identify putative novel links with BMI and obesity.

## Introduction

The rate of obesity is growing at an alarming rate worldwide − fast enough to call it an epidemic. As obesity is a risk factor for developing typically related diseases such as Type 2 Diabetes Mellitus (DM2), cardiovascular disease and some cancers (Wang *et al*. 2011), the situation is becoming a public health concern. The percentage of obesity is rising nationwide, with current adult obesity rates at close to 40%, up from 32% in 2004 (Ogden *et al*. 2006; TFAHRWJF 2018). In Nevada, the current adult obesity rate (BMI ≥ 30) is 27%, an increase from 21% in 2005 (TFAHRWJF 2018). Additionally, since 2016, Nevada witnessed a significant increase in the percentage of adults who are overweight (the current rate is 66%) (TFAHRWJF 2018). Studies identified several genetic factors that influence the development of obesity with estimates on the heritability of the disease (40%-75%) (Stunkard *et al*. 1986; 1990; Maes *et al*. 1997; Herrera and Lindgren 2010) and 65-80% (Malis *et al*. 2005).

High body mass index (BMI) and DM2 are known from many sources to be strongly related both epidemiologically and genetically (Kopelman 2007; Bays *et al*. 2007; Grarup *et al*. 2014; Cronin *et al*. 2014); however, these two conditions share very few known causative variants (Grarup *et al*. 2014; Karaderi *et al*. 2015). Although a number of large meta-analyses of multiple genome-wide association studies (GWASs) have detected possible causative single nucleotide polymorphisms (SNPs) of obesity and increased BMI (Scuteri *et al*. 2007; Frayling *et al*. 2007; Dina *et al*. 2007; Zeggini *et al*. 2007; Yanagiya *et al*. 2007; Hinney *et al*. 2007; Hunt *et al*. 2008; Price *et al*. 2008; Grant *et al*. 2008; Hotta *et al*. 2008; Loos *et al*. 2008; Tan *et al*. 2008; Villalobos-Comparán *et al*. 2008; Thorleifsson *et al*. 2008; Willer *et al*. 2009; Meyre *et al*. 2009; Wing *et al*. 2009; Liu *et al*. 2009; Shimaoka *et al*. 2010; Fawcett and Barroso 2010; Speliotes *et al*. 2010; Wang *et al*. 2011; Prakash *et al*. 2011; Okada *et al*. 2012; Cha *et al*. 2012; Berndt *et al*. 2013; Wheeler *et al*. 2013; Graff *et al*. 2013; Olza *et al*. 2013; Boender *et al*. 2014; Qureshi *et al*. 2017; Huđek *et al*. 2018; Gonzalez-Herrera *et al*. 2018), none, to the best of our knowledge, have included comprehensive GWASs on the quantitative BMI metric and on extreme obesity case-control simultaneously, as well as investigated phenotypic associations with BMI, obesity, and significant loci identified by the GWAS.

Our study begins with the Healthy Nevada Project (HNP), a project centered around a Northern Nevada cohort formed in 2016 and 2017 by Renown Health and the Desert Research Institute in Reno, NV to investigate factors that may contribute to health outcomes in Northern Nevada. Its first phase provided 10,000 individuals in Northern Nevada with genotyping using the 23andMe platform at no cost. Renown Health is the only tertiary care health system in the area, and 75% of these 10,000 individuals are cross-referenced in its extensive electronic health records (EHR) database. The Renown EHR database contains 86,610 BMI measurements for these 10,000 individuals over twelve years, along with comprehensive disease diagnoses, (e.g. diabetes or eating disorders) and other general conditions such as pregnancy, allowing for precise individual phenotypic classifications and thereby leading to more robust and meaningful phenotype-genotype associations.

The focus of the comprehensive GWAS-PheWAS examinations of the Healthy Nevada Project (HNP) cohort and its EHR database is two-fold: the first is to establish infrastructure to perform large-scale genome-wide and phenome-wide association investigations in alliance with complex electronic health care records; the second is to validate well-known published variants and associations with BMI and obesity in this cohort, as well as to identify possibly novel genotypic and phenotypic associations with BMI and extreme obesity.

The three GWASs identified several of the “usual suspects” for both BMI and obesity, such as *FTO* and *NEGR1*, that were shown to have a role in weight regulation (Scuteri *et al*. 2007; Frayling *et al*. 2007; Dina *et al*. 2007; Zeggini *et al*. 2007; Hinney *et al*. 2007; Hunt *et al*. 2008; Price *et al*. 2008; Grant *et al*. 2008; Hotta *et al*. 2008; Loos *et al*. 2008; Tan *et al*. 2008; Villalobos-Comparán *et al*. 2008; Thorleifsson *et al*. 2008; Willer *et al*. 2009; Meyre *et al*. 2009; Wing *et al*. 2009; Shimaoka *et al*. 2010; Fawcett and Barroso 2010; Speliotes *et al*. 2010; Herrera and Lindgren 2010; Wang *et al*. 2011; Prakash *et al*. 2011; Okada *et al*. 2012; Berndt *et al*. 2013; Wheeler *et al*. 2013; Graff *et al*. 2013; Olza *et al*. 2013; Boender *et al*. 2014; Qureshi *et al*. 2017; Gonzalez-Herrera *et al*. 2018). However, this study also identified a number of novel BMI and obesity associations to genes which are differentially expressed in obese patients (Jiao *et al*. 2008; Pietiläinen *et al*. 2008; Nakajima *et al*. 2016).

Additionally, using linked EHR, the PheWASs examined the pleiotropy of HNP BMI and obesity associated SNPs: whether these variants are linked with other endocrine or metabolic diagnoses or conditions of other nature. A second PheWAS identified many known phenotypes related to BMI and obesity, especially to DM2, abnormal glucose levels, hypertension, hyperlipidemia, sleep apnea, asthma and other less-studied BMI-related diagnoses.

## Materials and Methods

### The Renown EHR Database

The Renown Health EHR system was instated in 2007 on the EPIC system (EPIC System Corporation, Verona, Wisconsin, USA), and currently contains lab results, diagnosis codes (ICD9 and ICD10) and demographics of more than one million patients seen in the hospital system since 2005.

### Sample Collection

Saliva as a source of DNA was collected from 10,000 adults in Northern Nevada as the first phase of the Healthy Nevada Project to contribute to comprehensive population health studies in Nevada. The personal genetics company 23andMe, Inc. was used to genotype these individuals using the Oragene DX OGD-500.001 saliva kit [DNA Genotek, Ontario, Canada]. Genotypes are based on the Illumina Human OmniExpress-24 BeadChip platform [San Diego, CA, USA], that include approximately 570,000 SNPs.

### IRB and ethics statement

The study was reviewed and approved by the University of Nevada, Reno Institutional Review Board (IRB, project 956068-12). Participants in the Healthy Nevada Project undergo written and informed consent to having genetic information associated with electronic health information in a deidentified manner. All participants were eighteen years of age or older. Neither researchers nor participants have access to the complete EHR data and cannot map participants to patient identifiers. Patient identifiers are not incorporated into the EHR; rather, EHR and genetic data are linked in a separate environment via a unique identifier as approved by the IRB.

### Processing of EHR data

Most cohort participants had multiple BMI recordings across the thirteen years of EHR; the mean number of BMI records across the individuals was 12.2 records, with 215 the maximum number of records for the cohort. For the 5,811 individuals with more than one recorded BMI measure, a simple quality control step was first performed before computing the average BMI value. More specifically, the coefficient of variation (CV) across the multiple BMI records for each participant was computed; for those with CV in the 90^th^ quantile, any outlying BMI measure more than 1.5 standard deviations away from the mean BMI measure was excluded. Only 10% of participants’ computed coefficients of variation was less than 0.10, indicating little variation exists across multiple records in most individuals. The additional quality control step excluded one or more outlying BMI records in 106 individuals; these 701 BMI records included values such as “2823.42” and values less than 10. Examples of outliers include 158.38 in an individual’s set of values with mean 25.3 and “2874” in an individual with mean BMI measure of 22.4. Additionally, this quality control step allowed the study to include pregnant women: of the 464 pregnant women with BMI recorded for pregnant and non-pregnant phases, outlying pregnancy-related BMI records were easily identified and removed. The raw BMI values and quality-controlled average BMI values are presented in Supplementary Figure S1.

### Genotyping and Quality Control

Genotyping was performed by 23andMe using the Illumina Infimum DNA Human OmniExpress-24 BeadChip V4 (Illumina, San Diego, CA). This genotyping platform consists of approximately 570,000 SNPs. DNA extraction and genotyping were performed on saliva samples by the National Genetics Institute (NG1), a CLIA licensed clinical laboratory and a subsidiary of the Laboratory Corporation of America.

Raw genotype data were processed through a standard quality control process (Anderson *et al*. 2010; Verma *et al*. 2016; Schlauch *et al*. 2016; Verma *et al*. 2018; Schlauch *et al*. 2018). SNPs with a minor allele frequency (MAF) less than 0.005 were removed. SNPs that were out of Hardy Weinberg equilibrium (*p*-value < 1×10^-6^) were also excluded. Any SNP with a call rate less than 95% was removed; any individual with a call rate less than 95% was also excluded from further study. There was an observable bias within the African American sub-cohort, thus 89 African American participants were excluded from this study. Additionally, 107 patients with Type I Diabetes were removed, as were 29 participants with eating disorders recorded into their health record. After quality control, this left 500,508 high-quality SNPs and 6,645 participants in the BMI cohort with mean autosomal heterozygosity of 0.318. The same process yielded 5,994 participants when all individuals with DM2 diagnoses were removed. Within the extreme obesity study, participants with BMI values between and 25 were considered as controls, while any participant with BMI at least 35 kg/m^2^ was considered a case subject. Again, any individual with Type I Diabetes and recorded eating disorders was removed. This resulted in a cohort size of 2,994 participants with 984 extreme obese cases and 2,012 lean controls with a mean autosomal heterozygosity of 0.316.

A standard principal component analysis (PCA) was performed on the genotype data to identify principal components to correct for population substructure. Genotype data were pruned to exclude SNPs with high linkage disequilibrium using *PLINK* v1.9 (Purcell *et al*. 2007) and standard pruning parameters of 50 SNPs per sliding window; window size of five SNPs; *r*^2^=0.5 (Anderson *et al*. 2010). Regression models were adjusted by the first four principal components, decreasing the genomic inflation factor of all obesity and BMI traits to *λ* ≤ 1.06.

### Genome-Wide Association Studies

Using *PLINK*, we first performed a simple linear regression of BMI vs. genotype using the additive model (number of copies of the minor allele) including age, gender and the first four principal components as covariates to correct for any bias generated by these variables. In the first BMI study, participants with DM2 were excluded. The second BMI study included DM2-diagnosed participants and included DM2 as a covariate in the statistical model. To test associations between obesity and genotype, a standard case-control logistic regression was applied, adjusting for age, gender and the first four principal components. Total phenotypic variance explained by the SNPs was calculated by first producing a genetic relationship matrix of all SNPs on autosomal chromosomes in *PLINK*. Subsequently, a restricted maximum likelihood analysis was conducted using GTCA (Yang *et al*. 2011) on the relationship matrix to estimate the variance explained by the SNPs.

### Analysis of Variance

The mean BMI values across genotypes presented in Supplementary Tables S1 and S2 correlate with negative and positive effect sizes: SNPs showing a negative effect size have a decrease in mean BMI values across the genotypes from left to right (homozygous in major allele, heterozygous, homozygous in minor allele). The 6,645 log-transformed quality-controlled and averaged BMI measures were nearly normally distributed. As one-way ANOVA computations are robust against even moderate deviations of normality (Blanca *et al*. 2017), parametric ANOVA methods were used to make comparisons across the genotypes. All ANOVA F-test *p*-values of the significant SNPs identified in the two BMI studies are statistically significant at the alpha=0.05 level, even after a simple Bonferroni correction (.05/27 =0.0019, and .05/20=0.0025, respectively). Supplementary Table S3 presents the proportion of obese cases across each genotype. A simple test of equal proportions (Pearson’s chi-square test) is performed across these proportions. All *p*-values associated with the test of equal proportions in Supplementary Table S3 are also statistically significant at the 0.05 significance level upon a conservative Bonferroni multiple testing adjustment (.05/34=-.0015).

### Power of GWAS

The software program QUANTO (Gauderman 2002) was used to calculate sample sizes to detect effect sizes in the range [0.5,1] and odds ratios in [1,1.5] with at least 80% power under the additive model, at a two-sided Type I error level of 5%. Using the rate of extreme obesity (BMI ≥ 35 kg/m^2^) as 14.5% from Ogden et al. (Ogden *et al*. 2006), the case-control GWAS study of approximately 1,000 cases and 2,000 controls has sufficient statistical power (≥ 80%) with MAFs of 16% or greater to detect odds ratios of 1.225 or greater. As the MAF increases, the power to detect smaller odds ratios increases: for example, with a MAF of 25%, our sample size was adequate to detect odds ratios of size 1.18 or higher. With a small MAF of 8%, the power was also at least 80% to detect effect sizes as small as 0.58 in the BMI GWAS cohort of 6,645. With MAF of 17%, power was at least 80% to detect effect sizes as small as 0.425. Larger MAFs clearly can detect larger effect sizes with the same sample size. Specific effect sizes and MAFs can be seen in Table 2.

### Phenome-Wide Association Study

The **R** package PheWAS (Carroll *et al*. 2014) was used to perform two independent PheWAS analyses for each of our studies. The first examined associations between statistically significant SNPs identified in the respective GWASs and EHR phenotypes based on ICD codes. The second PheWAS identified associations between BMI levels or incidence of obesity, respectively, and ICD-based diagnoses. ICD9 and ICD10 codes for each individual in the cohort recorded in the Renown EHR were aggregated via a mapping from the Center for Medicare and Medicaid services (https://www.cms.gov/Medicare/Coding/ICD10/2018-ICD-10-CM-and-GEMs.html). A total of 22,693 individual diagnoses mapped to 4,769 documented ICD9 codes. ICD9 codes were aggregated and converted into 1,814 individual phenotype groups (“phecodes”) using the PheWAS package as described in Carroll and Denny (Denny *et al*. 2013; Carroll *et al*. 2014). Of these, only the phecodes that included at least 20 cases were used for downstream analyses, following Carroll’s protocol (Carroll *et al*. 2014). Age and gender were standard covariates included in the PheWAS models. The first type of PheWAS detected associations between statistically significant SNPs (*p*<1×10^-5^) identified in each of the three GWASs above and case/control status of EHR phenotypes represented by ICD codes. Specifically, a logistic regression between the incidence (number of cases) of each phenotype group (phecode) and the additive genotypes of each statistically significant SNP was performed, including age and gender as covariates. Possible associations of the phecodes with at least 20 individuals with each previously detected SNP were assessed. Two levels of significance were computed: the first, on which the reported results are based, was generated by first calculating the adjusted *p*-values for the multiple hypothesis tests using the Benjamini-Hochberg false discovery rate (FDR) (Benjamini and Hochberg 1995) and selecting the raw *p*-value corresponding to the FDR = 0.1 significance level, following Denny’s protocol (Denny *et al*. 2013). This level is represented by a red line in PheWAS images. The second, more conservative, significance level was computed as a Bonferroni correction for all possible associations made in this analysis: *p*=0.05 / *N_ps_*, where *N_ps_* is the sum of the number of phecodes tested for each individual SNP, across all identified SNPs. This significance level is represented by a blue line in PheWAS images.

A second PheWAS, as outlined in Carroll et al. (2014) (Carroll *et al*. 2014), was performed to examine associations between BMI, and secondarily, obesity, and the phecodes. Specifically, a linear regression between the BMI measures and the case/control status of a phecode was performed (with age and gender as covariates) for each phecode including at least 20 individuals. Significance levels corresponded to the FDR value of 0.1 and are not shown in either figure due to space constraints. The Bonferroni corrections for the BMI study and obesity study were 3.3×10^-5^ and 3.7×10^-5^, respectively, notably less conservative than the 1×10^-15^ significance level represented in the images.

## Results

### Characteristics of cohort

Our study consisted of 6,870 genotyped participants which had measures for age at consent, gender, ethnicity and BMI. From those genotyped individuals, we removed 107 participants who had Type I Diabetes, as well as 29 individuals who had any type of eating disorder. Preliminary quality control of the genotype data demonstrated a strong genetic bias within the African American subpopulation, and thus 89 African American participants were excluded prior to association analysis. Our final cohort characteristics of 6,645 individuals are described in Table 1, which illustrates the makeup of the cohort with respect to gender, age, ethnic origin, and standardized value of BMI after removal of outliers using a custom algorithm. For the extreme obesity (Class 2 and 3) case vs. control study, the normal (healthy) control range consisted of BMI values between 18.5 and 25 kg/m^2^, while the case obese values were any BMI ≥ 35 kg/m^2^ (Hruby and Hu 2014). The number of participants in each range is displayed in column two of Table 1.

**Table 1.** Cohort Characteristics.

|  | Association with BMI Measures | Association with Obesity |
| --- | --- | --- |
| Cohort Size | 6,645 | 2,996 |
| Age (years) | 50.91 ± 15.97 | 48.91 ± 15.96 |
| Male (%) | 2145 (32.28) | 747 (24.93) |
| African American (%) | 0 (0) | 0 (0) |
| Asian (%) | 157 (2.36) | 84 (2.80) |
| Caucasian (%) | 5,945 (89.47) | 2653 (88.55) |
| Latino (%) | 173 (2.60) | 81 (2.70) |
| Native American (%) | 40 (0.60) | 18 (0.60) |
| Pacific Islander (%) | 13 (0.20) | 4 (0.13) |
| Unknown (%) | 317 (4.77) | 156 (5.21) |
| DM2 | 651 (9.8) | 246 (8.2) |
| Quality-Controlled BMI Range | 28.58 ± 6.41 | 22.57 ± 1.64 (Control) |
|  |  | 40.15 ± 4.99 (Cases) |

Table of cohort characteristics. Continuous variables are presented as mean ± SD; categorical variables are presented as counts and percentages. All values were standardized to using a custom algorithm to remove outliers. BMI has the units of kg/m<sup>2</sup>.

### GWAS of BMI in the Healthy Nevada Cohort

Using quality-controlled BMI values, two separate GWASs were performed to find genotypic associations with BMI using *PLINK*. In the first association study, all individuals diagnosed with Type 2 Diabetes (DM2) were excluded to focus on the association between the genotype and BMI under the additive model with adjustments for gender, age and the first four principal components (PC1-PC4). The second association analysis included all DM2-diagnosed individuals and added DM2 as a covariate to the model, again using the additive model with adjustments for gender, age, diabetes status, and PC1-PC4. Genomic inflation coefficients (lambda) were computed for the two separate cohorts: 1.06 for the association without DM as a covariate, and 1.06 for the association where DM is a covariate. Any SNP with association *p*-value of *p* < 1×10^-5^ was considered a statistically significant association, based on the standard of the NHGRI-EBI Catalog of published genome-wide association studies [https://www.ebi.ac.uk/gwas/docs/methods/criteria], as well as obesity studies performed by Frayling et al. (Frayling *et al*. 2007). Genetic variance in the BMI study with DM2 cases removed was 15.78%; genetic variance was 17.49% in the BMI study with DM2 cases included.

The first GWAS was performed on 5,994 total participants without DM2 and identified 20 SNPs across seven chromosomes at statistical significance defined by *p*<1×10^-5^ (Table 2). The majority of these mapped to the *FTO* gene on chromosome 16, while two SNPs mapped to *TDH* on chromosome 8 (Supplementary Figure S2). Of the 20 SNPs, 15 were shown to be associated with BMI in previous publications (Scuteri *et al*. 2007; Frayling *et al*. 2007; Dina *et al*. 2007; Zeggini *et al*. 2007; Yanagiya *et al*. 2007; Hinney *et al*. 2007; Hunt *et al*. 2008; Price *et al*. 2008; Grant *et al*. 2008; Hotta *et al*. 2008; Loos *et al*. 2008; Tan *et al*. 2008; Villalobos-Comparán *et al*. 2008; Thorleifsson *et al*. 2008; Willer *et al*. 2009; Meyre *et al*. 2009; Wing *et al*. 2009; Liu *et al*. 2009; Shimaoka *et al*. 2010; Fawcett and Barroso 2010; Speliotes *et al*. 2010; Wang *et al*. 2011; Prakash *et al*. 2011; Okada *et al*. 2012; Cha *et al*. 2012; Berndt *et al*. 2013; Wheeler *et al*. 2013; Graff *et al*. 2013; Olza *et al*. 2013; Boender *et al*. 2014; Qureshi *et al*. 2017; Huđek *et al*. 2018; Gonzalez-Herrera *et al*. 2018). A large majority of the SNPs (17/20) lie within noncoding regions of genes and are intronic in nature. It is interesting to note that our strongest associations lie within the *FTO* gene (*p* < 3.5×10^-6^). Results are presented in Table 2: **BMI without DM2** lists the significant associations of our cohort that exclude all DM2 diagnoses. **BMI with DM2** presents significant associations with BMI in the cohort that includes participants with a DM2 diagnosis. Effect sizes (and their standard deviations) are presented as change in BMI per each copy of the minor allele. Raw per-SNP *p*-values are presented.

**Table 2.**
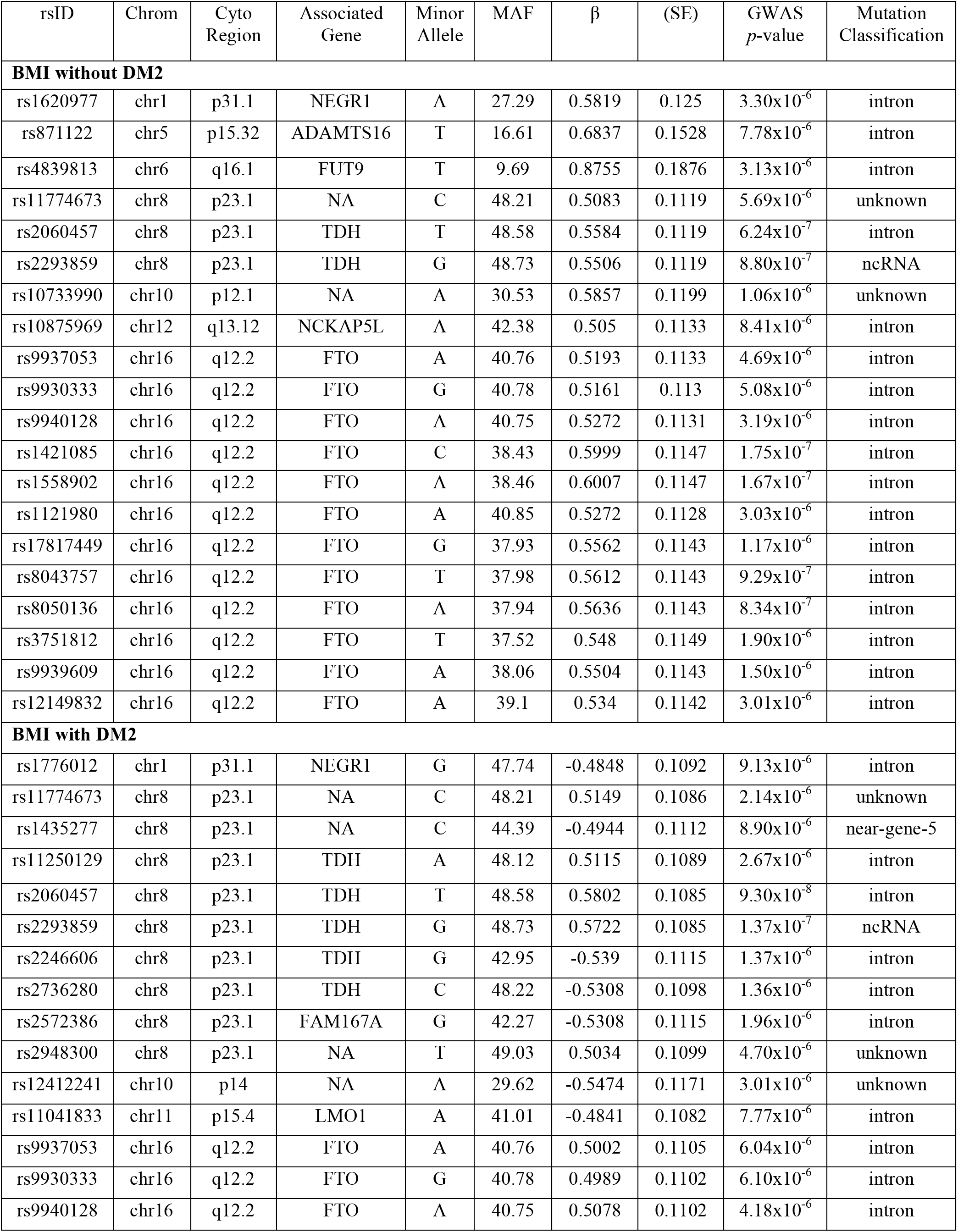
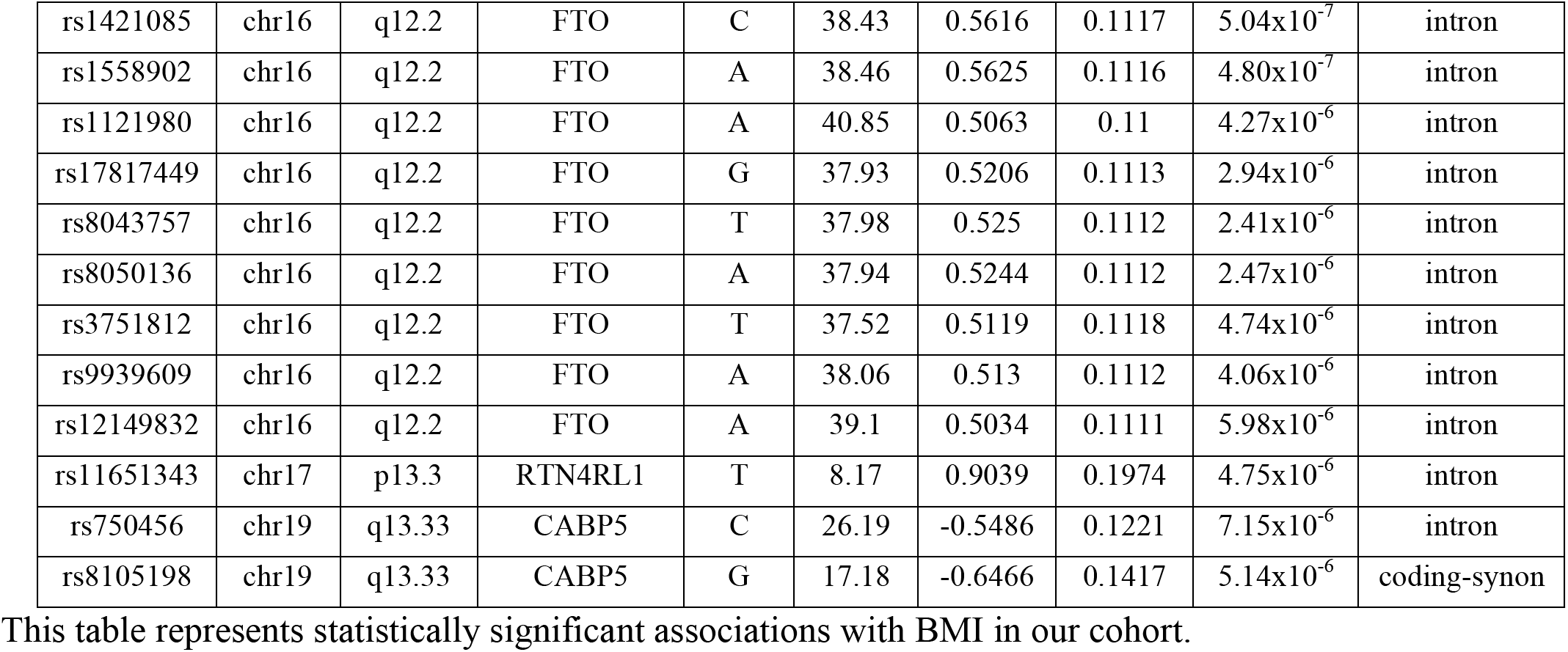
Statistically Significant BMI GWAS SNPs.

| rsID | Chrom | Cyto Region | Associated Gene | Minor Allele | MAF | $\beta$ | (SE) | GWAS *p*-value | Mutation Classification |
| --- | --- | --- | --- | --- | --- | --- | --- | --- | --- |
| **BMI without DM2** |  |  |  |  |  |  |  |  |  |
| rs1620977 | chr1 | p31.1 | NEGR1 | A | 27.29 | 0.5819 | 0.125 | 3.30x10<sup>-6</sup> | intron |
| rs871122 | chr5 | p15.32 | ADAMTS16 | T | 16.61 | 0.6837 | 0.1528 | 7.78x10<sup>-6</sup> | intron |
| rs4839813 | chr6 | q16.1 | FUT9 | T | 9.69 | 0.8755 | 0.1876 | 3.13x10<sup>-6</sup> | intron |
| rs11774673 | chr8 | p23.1 | NA | C | 48.21 | 0.5083 | 0.1119 | 5.69x10<sup>-6</sup> | unknown |
| rs2060457 | chr8 | p23.1 | TDH | T | 48.58 | 0.5584 | 0.1119 | 6.24x10<sup>-7</sup> | intron |
| rs2293859 | chr8 | p23.1 | TDH | G | 48.73 | 0.5506 | 0.1119 | 8.80x10<sup>-7</sup> | ncRNA |
| rs10733990 | chr10 | p12.1 | NA | A | 30.53 | 0.5857 | 0.1199 | 1.06x10<sup>-6</sup> | unknown |
| rs10875969 | chr12 | q13.12 | NCKAP5L | A | 42.38 | 0.505 | 0.1133 | 8.41x10<sup>-6</sup> | intron |
| rs9937053 | chr16 | q12.2 | FTO | A | 40.76 | 0.5193 | 0.1133 | 4.69x10<sup>-6</sup> | intron |
| rs9930333 | chr16 | q12.2 | FTO | G | 40.78 | 0.5161 | 0.113 | 5.08x10<sup>-6</sup> | intron |
| rs9940128 | chr16 | q12.2 | FTO | A | 40.75 | 0.5272 | 0.1131 | 3.19x10<sup>-6</sup> | intron |
| rs1421085 | chr16 | q12.2 | FTO | C | 38.43 | 0.5999 | 0.1147 | 1.75x10<sup>-7</sup> | intron |
| rs1558902 | chr16 | q12.2 | FTO | A | 38.46 | 0.6007 | 0.1147 | 1.67x10<sup>-7</sup> | intron |
| rs1121980 | chr16 | q12.2 | FTO | A | 40.85 | 0.5272 | 0.1128 | 3.03x10<sup>-6</sup> | intron |
| rs17817449 | chr16 | q12.2 | FTO | G | 37.93 | 0.5562 | 0.1143 | 1.17x10<sup>-6</sup> | intron |
| rs8043757 | chr16 | q12.2 | FTO | T | 37.98 | 0.5612 | 0.1143 | 9.29x10<sup>-7</sup> | intron |
| rs8050136 | chr16 | q12.2 | FTO | A | 37.94 | 0.5636 | 0.1143 | 8.34x10<sup>-7</sup> | intron |
| rs3751812 | chr16 | q12.2 | FTO | T | 37.52 | 0.548 | 0.1149 | 1.90x10<sup>-6</sup> | intron |
| rs9939609 | chr16 | q12.2 | FTO | A | 38.06 | 0.5504 | 0.1143 | 1.50x10<sup>-6</sup> | intron |
| rs12149832 | chr16 | q12.2 | FTO | A | 39.1 | 0.534 | 0.1142 | 3.01x10<sup>-6</sup> | intron |
| **BMI with DM2** |  |  |  |  |  |  |  |  |  |
| rs1776012 | chr1 | p31.1 | NEGR1 | G | 47.74 | -0.4848 | 0.1092 | 9.13x10<sup>-6</sup> | intron |
| rs11774673 | chr8 | p23.1 | NA | C | 48.21 | 0.5149 | 0.1086 | 2.14x10<sup>-6</sup> | unknown |
| rs1435277 | chr8 | p23.1 | NA | C | 44.39 | -0.4944 | 0.1112 | 8.90x10<sup>-6</sup> | near-gene-5 |
| rs11250129 | chr8 | p23.1 | TDH | A | 48.12 | 0.5115 | 0.1089 | 2.67x10<sup>-6</sup> | intron |
| rs2060457 | chr8 | p23.1 | TDH | T | 48.58 | 0.5802 | 0.1085 | 9.30x10<sup>-8</sup> | intron |
| rs2293859 | chr8 | p23.1 | TDH | G | 48.73 | 0.5722 | 0.1085 | 1.37x10<sup>-7</sup> | ncRNA |
| rs2246606 | chr8 | p23.1 | TDH | G | 42.95 | -0.539 | 0.1115 | 1.37x10<sup>-6</sup> | intron |
| rs2736280 | chr8 | p23.1 | TDH | C | 48.22 | -0.5308 | 0.1098 | 1.36x10<sup>-6</sup> | intron |
| rs2572386 | chr8 | p23.1 | FAM167A | G | 42.27 | -0.5308 | 0.1115 | 1.96x10<sup>-6</sup> | intron |
| rs2948300 | chr8 | p23.1 | NA | T | 49.03 | 0.5034 | 0.1099 | 4.70x10<sup>-6</sup> | unknown |
| rs12412241 | chr10 | p14 | NA | A | 29.62 | -0.5474 | 0.1171 | 3.01x10<sup>-6</sup> | unknown |
| rs11041833 | chr11 | p15.4 | LMO1 | A | 41.01 | -0.4841 | 0.1082 | 7.77x10<sup>-6</sup> | intron |
| rs9937053 | chr16 | q12.2 | FTO | A | 40.76 | 0.5002 | 0.1105 | 6.04x10<sup>-6</sup> | intron |
| rs9930333 | chr16 | q12.2 | FTO | G | 40.78 | 0.4989 | 0.1102 | 6.10x10<sup>-6</sup> | intron |
| rs9940128 | chr16 | q12.2 | FTO | A | 40.75 | 0.5078 | 0.1102 | 4.18x10<sup>-6</sup> | intron |
| rs1421085 | chr16 | q12.2 | FTO | C | 38.43 | 0.5616 | 0.1117 | $5.04 \times 10^{-7}$ | intron |
| rs1558902 | chr16 | q12.2 | FTO | A | 38.46 | 0.5625 | 0.1116 | $4.80 \times 10^{-7}$ | intron |
| rs1121980 | chr16 | q12.2 | FTO | A | 40.85 | 0.5063 | 0.11 | $4.27 \times 10^{-6}$ | intron |
| rs17817449 | chr16 | q12.2 | FTO | G | 37.93 | 0.5206 | 0.1113 | $2.94 \times 10^{-6}$ | intron |
| rs8043757 | chr16 | q12.2 | FTO | T | 37.98 | 0.525 | 0.1112 | $2.41 \times 10^{-6}$ | intron |
| rs8050136 | chr16 | q12.2 | FTO | A | 37.94 | 0.5244 | 0.1112 | $2.47 \times 10^{-6}$ | intron |
| rs3751812 | chr16 | q12.2 | FTO | T | 37.52 | 0.5119 | 0.1118 | $4.74 \times 10^{-6}$ | intron |
| rs9939609 | chr16 | q12.2 | FTO | A | 38.06 | 0.513 | 0.1112 | $4.06 \times 10^{-6}$ | intron |
| rs12149832 | chr16 | q12.2 | FTO | A | 39.1 | 0.5034 | 0.1111 | $5.98 \times 10^{-6}$ | intron |
| rs11651343 | chr17 | p13.3 | RTN4RL1 | T | 8.17 | 0.9039 | 0.1974 | $4.75 \times 10^{-6}$ | intron |
| rs750456 | chr19 | q13.33 | CABP5 | C | 26.19 | -0.5486 | 0.1221 | $7.15 \times 10^{-6}$ | intron |
| rs8105198 | chr19 | q13.33 | CABP5 | G | 17.18 | -0.6466 | 0.1417 | $5.14 \times 10^{-6}$ | coding-synon |

This table represents statistically significant associations with BMI in our cohort.

Statistical analysis with *PLINK* demonstrated that DM2 is a significant predictor of BMI, with the *p*-value of its coefficient consistently less than *p* < 2×10^-16^ in each per-SNP linear regression. The entire cohort includes 6,645 participants: of those, 651 have a diagnosis of DM2 in their twelve-year medical history. A GWAS applied to this larger cohort identified 27 statistically significant SNPs across seven chromosomes associated with BMI at *p*<1×10^-5^ (Figure 1). In comparison to the original GWAS (without DM2 individuals), 75% of the SNPs (15/20) were also found to be associated with BMI in this association. Furthermore, 77% of the SNPs in this second GWAS (21/27) were previously associated with BMI in previous research studies (Scuteri *et al*. 2007; Frayling *et al*. 2007; Dina *et al*. 2007; Zeggini *et al*. 2007; Yanagiya *et al*. 2007; Hinney *et al*. 2007; Hunt *et al*. 2008; Price *et al*. 2008; Grant *et al*. 2008; Hotta *et al*. 2008; Loos *et al*. 2008; Tan *et al*. 2008; Villalobos-Comparán *et al*. 2008; Thorleifsson *et al*. 2008; Willer *et al*. 2009; Meyre *et al*. 2009; Wing *et al*. 2009; Liu *et al*. 2009; Shimaoka *et al*. 2010; Fawcett and Barroso 2010; Speliotes *et al*. 2010; Wang *et al*. 2011; Prakash *et al*. 2011; Okada *et al*. 2012; Cha *et al*. 2012; Berndt *et al*. 2013; Wheeler *et al*. 2013; Graff *et al*. 2013; Olza *et al*. 2013; Boender *et al*. 2014; Christensen *et al*. 2015; Nakajima *et al*. 2016; Thomsen *et al*. 2016; Qureshi *et al*. 2017; Huđek *et al*. 2018; Gonzalez-Herrera *et al*. 2018). With the addition of DM2 as a covariate, the GWAS identified several additional SNPs on chromosome 8, as well as SNPs on chromosomes 17 and 19. These additional SNPs were previously linked to BMI and obesity in other studies (Christensen *et al*. 2015; Nakajima *et al*. 2016; Thomsen *et al*. 2016). Manhattan plots for the two BMI GWAS studies are presented in Figure 1 and Supplementary Figure S2, with the linear associations results presented in Table 2.

**Figure 1:**
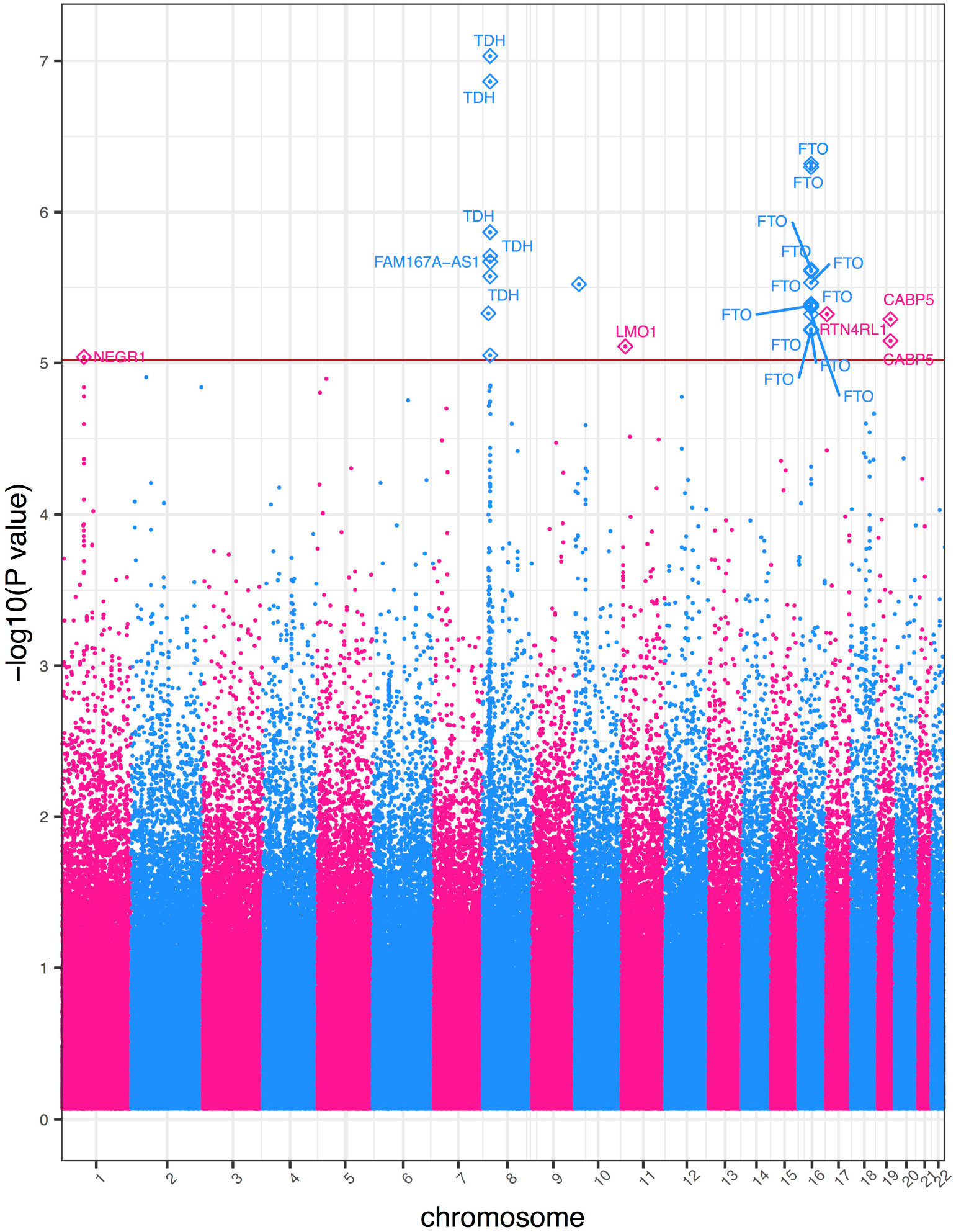
Manhattan plot of GWAS results of BMI including DM2-diagnosed individuals. Genome-wide association study results for BMI. This study includes DM2-diagnosed individuals and the statistical model includes DM2 as a bimodal covariate. The *x*-axis represents the genomic position of 500,508 SNPs. The *y*-axis represents -log_10_-transformed raw *p*-values of each genotypic association. The red horizontal line indicates the significance level 1×10^-5^.

The SNP on chromosome 17 is of particular interest, as it has the largest effect of any SNP identified in our study (β=0.90). It is also the rarest SNP tested in our cohort with minor allele frequency (MAF) 8.17%. The median MAF across the strongest associative SNPs in both studies is 40%, which demonstrates that most of the SNPs are common and thus result in relatively moderate individual effect sizes. Most of the SNPs lie within noncoding intronic regions. While these SNPs would not alter the amino acid coding sequence of the translated protein, several previous studies articulated that polymorphisms within introns can affect intron splicing as well as transcriptional and translational efficiency, and therefore may be linked to disease (Lalonde *et al*. 2011).

### Case-Control GWAS of Extreme Obesity in the Healthy Nevada Cohort

A complementary GWAS was performed to identify genotype-phenotype links in extreme obesity (BMI ≥ 35) versus non-obese (BMI between 18.5 and 25 kg/m^2^) in our cohort. This study incorporated 2996 participants (984 extreme obese cases, 2012 non-obese controls), and under the log-additive model with adjustments for gender, age and the first four principal components, identified 26 SNPs across six chromosomes that were associated with obesity at *p* < 1×10^-5^, with approximately 70% associated with obesity and BMI in prior studies (Figure 2). The percentage of phenotypic variance attributed to genetic variation was 15.7%. The genomic inflation coefficient (lambda) for the obesity cohort was computed as 1.05. We also include eight SNPs found slightly above the significance threshold in the *FTO* gene that are reported in several studies as obesity-related (Ehrlich and Friedenberg 2016; West *et al*. 2018). In comparison to the two quantitative-trait BMI GWASs, this study identified several more associations around the *NEGR1* gene on chromosome 1. We also identified SNPs in two genes, *PFKFB3* and *CABP5*, which are associated with obesity in other studies (Scuteri *et al*. 2007; Jiao *et al*. 2008; Nakajima *et al*. 2016). Note that all the mutations in the *FTO* gene increase the odds of obesity risk. Table 3 lists the strongest SNPs associated in our extreme obese vs. non-obese GWAS. Effect sizes and their standard deviations are presented as odds ratios. Raw *p*-values generated by the GWAS are also presented.

**Figure 2:**
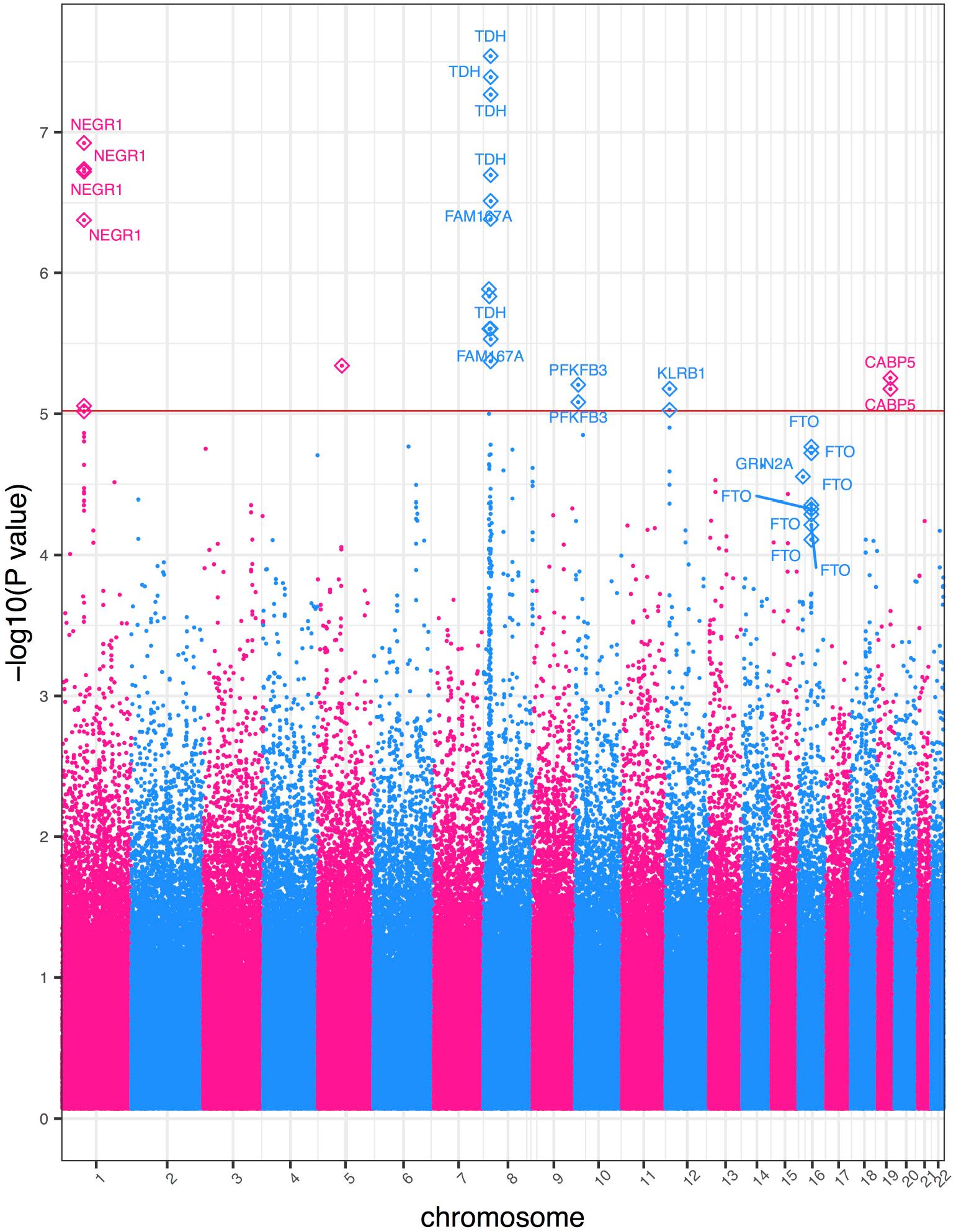
Obesity Case-Control GWAS Manhattan Plot. Genome-wide association study results for the case-control obesity study. This cohort includes DM2-diagnosed individuals. The *x*-axis represents the genomic position of 500,508 SNPs. The *y*-axis represents -log_10_-transformed raw *p*-values of each genotypic association. The red horizontal line indicates the significance level 1×10^-5^.

**Table 3.** Statistically Significant Obesity GWAS SNPs.

| rsID | Chrom | Cyto<br>Region | Associated<br>Gene | Minor<br>Allele | MAF | Odds<br>Ratio | (SE) | GWAS<br>*p*-value | Mutation<br>Classification |
| --- | --- | --- | --- | --- | --- | --- | --- | --- | --- |
| rs1776012 | chr1 | p31.1 | *NEGR1* | G | 47.36 | 0.7365 | 0.05775 | 1.19x10<sup>-7</sup> | intron |
| rs9424977 | chr1 | p31.1 | *NEGR1* | C | 47.09 | 0.7411 | 0.05745 | 1.83x10<sup>-7</sup> | intron |
| rs1620977 | chr1 | p31.1 | *NEGR1* | A | 27.31 | 1.369 | 0.06213 | 4.21x10<sup>-7</sup> | intron |
| rs1870676 | chr1 | p31.1 | *NEGR1* | T | 46.97 | 0.7403 | 0.05773 | 1.90x10<sup>-7</sup> | intron |
| rs3101336 | chr1 | p31.1 | NA | T | 35.93 | 0.7678 | 0.0597 | 9.60x10<sup>-6</sup> | unknown |
| rs2568958 | chr1 | p31.1 | NA | G | 35.94 | 0.767 | 0.05967 | 8.79x10<sup>-6</sup> | unknown |
| rs2815752 | chr1 | p31.1 | NA | G | 35.94 | 0.767 | 0.05967 | 8.79x10<sup>-6</sup> | unknown |
| rs2173676 | chr5 | q14.1 | NA | C | 26.52 | 0.7433 | 0.0647 | 4.56x10<sup>-6</sup> | unknown |
| rs11774673 | chr8 | p23.1 | NA | C | 48.25 | 1.304 | 0.05778 | 4.25x10<sup>-6</sup> | unknown |
| rs1435277 | chr8 | p23.1 | NA | C | 44.28 | 0.7419 | 0.05898 | 4.15x10<sup>-7</sup> | near-gene-5 |
| rs11250129 | chr8 | p23.1 | *TDH* | A | 48.26 | 1.314 | 0.05796 | 2.49x10<sup>-6</sup> | intron |
| rs2060457 | chr8 | p23.1 | *TDH* | T | 48.76 | 1.379 | 0.05797 | 2.88x10<sup>-8</sup> | intron |
| rs2293859 | chr8 | p23.1 | *TDH* | G | 48.81 | 1.375 | 0.05798 | 4.06x10<sup>-8</sup> | ncRNA |
| rs2246606 | chr8 | p23.1 | *TDH* | G | 42.76 | 0.7357 | 0.05905 | 2.01x10<sup>-7</sup> | intron |
| rs2736280 | chr8 | p23.1 | *TDH* | C | 48.23 | 0.727 | 0.05863 | 5.41x10<sup>-8</sup> | intron |
| rs2572386 | chr8 | p23.1 | *FAM167A* | G | 41.91 | 0.7378 | 0.05941 | 3.07x10<sup>-7</sup> | intron |
| rs1435282 | chr8 | p23.1 | *FAM167A* | A | 45.57 | 0.7631 | 0.05786 | 2.95x10<sup>-6</sup> | intron |
| rs2948300 | chr8 | p23.1 | NA | T | 49.06 | 1.325 | 0.05809 | 1.30x10<sup>-6</sup> | unknown |
| rs2953802 | chr8 | p23.1 | NA | A | 42 | 1.318 | 0.05734 | 1.47x10<sup>-6</sup> | unknown |
| rs435581 | chr8 | p23.1 | NA | A | 45.19 | 0.7609 | 0.05802 | 2.50x10<sup>-6</sup> | unknown |
| rs680951 | chr10 | p15.1 | *PFKFB3* | G | 33.83 | 0.7614 | 0.06116 | 8.26x10<sup>-6</sup> | intron |
| rs666595 | chr10 | p15.1 | *PFKFB3* | A | 31.47 | 0.7537 | 0.06256 | 6.22x10<sup>-6</sup> | intron |
| rs2058426 | chr12 | p13.31 | NA | A | 49.67 | 0.7798 | 0.05614 | 9.40x10<sup>-6</sup> | near-gene-3 |
| rs2241005 | chr12 | p13.31 | *KLRB1* | C | 49.98 | 0.7768 | 0.05608 | 6.65x10<sup>-6</sup> | intron |
| rs1421085 | chr16 | q12.2 | *FTO* | C | 38.18 | 1.28 | 0.05766 | 1.90x10<sup>-5</sup> | intron |
| rs1558902 | chr16 | q12.2 | *FTO* | A | 38.21 | 1.281 | 0.0576 | 1.72x10<sup>-5</sup> | intron |
| rs17817449 | chr16 | q12.2 | *FTO* | G | 37.63 | 1.261 | 0.05732 | 5.15x10<sup>-5</sup> | intron |
| rs8043757 | chr16 | q12.2 | *FTO* | T | 37.66 | 1.264 | 0.05733 | 4.42x10<sup>-5</sup> | intron |
| rs8050136 | chr16 | q12.2 | *FTO* | A | 37.66 | 1.263 | 0.05733 | 4.72x10<sup>-5</sup> | intron |
| rs3751812 | chr16 | q12.2 | *FTO* | T | 37.27 | 1.26 | 0.05764 | 6.13x10<sup>-5</sup> | intron |
| rs9939609 | chr16 | q12.2 | *FTO* | A | 37.83 | 1.254 | 0.05737 | 7.79x10<sup>-5</sup> | intron |
| rs2267770 | chr16 | p13.2 | *GRIN2A* | T | 41.01 | 0.7859 | 0.0575 | 2.79x10<sup>-5</sup> | intron |
| rs750456 | chr19 | q13.33 | *CABP5* | C | 26.1 | 0.7405 | 0.06615 | 5.58x10<sup>-6</sup> | intron |
| rs8105198 | chr19 | q13.33 | *CABP5* | G | 16.77 | 0.7026 | 0.07836 | 6.67x10<sup>-6</sup> | coding-synon |

This table presents statistically significant associations with extreme obesity in the case-control study. From our three separate GWASs, we identified fifteen different chromosomal cytoband regions across ten chromosomes associated with at least one BMI or obesity-related trait. All but three of those cytoband regions contained a gene, while the remaining cytoband hits were in noncoding regions of the genome. Approximately 70% of the SNPs identified in this study were linked to BMI and obesity in prior studies (Tables 2 and 3), validating our methods (Scuteri *et al*. 2007; Frayling *et al*. 2007; Dina *et al*. 2007; Yanagiya *et al*. 2007; Hinney *et al*. 2007; Jiao *et al*. 2008; Pietiläinen *et al*. 2008; Grant *et al*. 2008; Hotta *et al*. 2008; Thorleifsson *et al*. 2008; Joe *et al*. 2009; Nakajima *et al*. 2016; Thomsen *et al*. 2016; Justice *et al*. 2017). The functions of the genes which lie within the cytoband regions are outlined in Table 4.

**Table 4.** Table Presenting Gene Functions.

| Gene | Gene Description | Region | Trait | Function | Reference |
| --- | --- | --- | --- | --- | --- |
| NEGR1 | Neuronal Growth Regulator 1 | p31.1 | BMI No DM2<br>BMI w DM2 obesity | Cell-adhesion molecule and regulator of cellular processes as neurite outgrowth and synapse formation | (Hashimoto *et al.* 2008; Boender *et al.* 2014) |
| ADAMTS16 | ADAM Metallopeptidase with Thrombospondin Type 1 Motif 16 | p15.32 | BMI No DM2 | May be a secreted proteinase | (Surridge *et al.* 2009) |
| FUT9 | Fucosyltransferase 9 | q16.1 | BMI No DM2 | Plays a role in the biosynthesis of Lewis X (LeX) antigen precursor polysaccharides | (Gouveia *et al.* 2012) |
| TDH | L-Threonine Dehydrogenase | p23.1 | BMI No DM2<br>BMI w DM2 obesity | Likely pseudogene that cannot produce functional protein | (Yanagiya *et al.* 2007) |
| NCKAP5L | NCK Associated Protein 5 Like | q13.12 | BMI No DM2 | Regulates microtubule organization and stabilization | (Mori *et al.* 2015) |
| FTO | Fat Mass and Obesity Associated | q12.2 | BMI No DM2<br>BMI w DM2 obesity | Mediates oxidative demethylation of different RNA species. Acts as a regulator of fat and energy homeostasis | (Frayling *et al.* 2007; Jia *et al.* 2011; Wei *et al.* 2018) |
| FAM167A | Family with Sequence Similarity 167 Member A | p23.1 | BMI w DM2 obesity | Unknown function. May have a role in autoimmune diseases | (Chen *et al.* 2015) |
| LMO1 | LIM domain only 1 | p15.4 | BMI w DM2 | Transcriptional regulator through binding of transcription factors | (Wang *et al.* 2010) |
| RTN4RL1 | Reticulon 4 Receptor Like 1 | p13.3 | BMI w DM2 | Cell surface receptor | (Pignot *et al.* 2003) |
| CABP5 | Calcium Binding Protein 5 | q13.33 | BMI w DM2 obesity | Plays a role in calcium mediated cellular signal transduction | (Haeseleer *et al.* 2002) |
| PFKFB3 | 6-Phosphofructo-2-kinase/Fructose-2,6-biphosphatase 3 | p15.1 | obesity | Potent regulator of glycolysis | (Nelson and Cox 2005; De Bock *et al.* 2013) |
| KLRB1 | Killer Cell Lectin-like Receptor Subfamily B, Member 1 | p13.31 | obesity | Possible regular of natural killer cells | (Rother *et al.* 2015) |
| GRIN2A | Glutamate Ionotropic Receptor NMDA Type Subunit 2A | p13.2 | obesity | Possibly has a role in learning and long-term memory | (Micu *et al.* 2006) |

This table presents functions of genes associated to all SNPs found significantly associated to one or more BMI and obesity traits in the GWASs.

### Analysis of Variance

The mean BMI values across genotypes presented in Supplementary Tables S1 and S2 correlate with negative and positive effect sizes: SNPs showing a negative effect size have a decrease in mean BMI values across the genotypes from left to right (homozygous in major allele, heterozygous, homozygous in minor allele). Note that BMI levels increase with the increase of the number of minor alleles, which is typical of variants in *FTO* (Frayling *et al*. 2007). All ANOVA F-test *p*-values of the significant SNPs identified in the two BMI studies are statistically significant at the alpha=0.05 level, even after a simple Bonferroni correction (.05/27 =0.0019, and .05/20=0.0025, respectively). Supplementary Table S3 presents the proportion of extremely obese cases across each genotype. A box and whisker figure of ANOVA results for one of the strongest associations (rs9939609) is shown in Supplementary Figure S3.

### PheWAS of BMI and Obesity

Beyond the GWASs, we present here two comprehensive PheWAS studies that follow each GWAS. The first examines pleiotropy, i.e., whether additional phenotypic associations exist between the statistically significant SNPs associated with BMI or obesity in our cohort. The second investigates which EHR phenotype groups are associated with BMI; more specifically, the analysis identifies whether the number of individuals in an EHR phenotype group is a predictor of BMI and/or extreme obesity.

A PheWAS tested the 20 statistically significant SNPs identified in the first BMI GWAS (the sub-cohort with no DM2-diagnoses) for association with 562 EHR phenotypes and resulted in no statistically significant associations at the false discovery rate of 0.1 (data not shown). The top two associations showed that a locus on *FUT9* (rs4839813) associated with obesity, and rs1620977 on *NEGR1* associated with morbid obesity with raw *p*-values *p*=2.3×10^-5^ and 2.8×10^-5^, respectively. The second PheWAS identified 179 EHR (phenotypic) associations of BMI (DM2-diagnosed participants excluded) with *p*<2×10^-2^, that associates to an adjusted *p*-value of 0.1 (see Materials and Methods). Included in the strongest phenotypic associations are obesity, morbid obesity, and overweight (*p*<1×10^-80^), sleep apnea (*p*<1×10^-45^), hypertension (*p*<1×10^-40^), abnormal glucose (*p*<1×10^-25^), hyperlipidemia, asthma, GERD, osteoporosis, and others. (Data not shown).

The PheWAS of the second BMI GWAS (DM2-diagnosed individuals included) examined whether 633 EHR phenotype groups containing at least 20 participants are dependent on the genotypes of the 27 statistically significant SNPs associated with BMI in our cohort (Figure 3). Results of this PheWAS indicate that *TDH* and *FAMA167-AS1* are strongly associated with DM2. Variants in the *FTO* gene associate with obesity and overweight phenotypes. An association with hypertension and essential hypertension and the locus rs12412241 on chromosome 10 was detected (*p*<1×10^-4^). Several strong associations with *FTO* loci and the prostate-specific antigen (PSA) were also found. Variants in the *FTO* gene also associated with obesity at the *p*=3×10^-4^ level, and with hypercholesterolemia with *p*=1×10^-3^ level. Significant associations (*p*<1.02×10^-4^, associated to an adjusted *p*-value of 0.1, see Materials and Methods) are included in Table 5 and illustrated in Figure 3, where the blue line represents the Bonferroni correction of *p*=3×10^-6^. The second PheWAS examined links between BMI and the 1,523 EHR phenotype groups containing at least 20 individuals in this cohort and showed that 301 such clinical phenotypes groups associated with significance *p*<1.96×10^-2^. (This significance level is associated to an adjusted *p*-value of 0.1, as described in the Materials and Methods) These are shown in Figure 4. Included in the highest associations are obesity, morbid obesity, and overweight (*p*<1×10^-100^), DM2 (*p*<1×10^-87^), hypertension ((*p*<1×10^-82^), sleep apnea (*p*<1×10^-80^), abnormal glucose (*p*<1×10^-53^), hyperlipidemia, asthma and other respiratory disorders, GERD, edema, liver disease, mood disorders, polycystic ovaries, and others. Significant associations are presented in Supplementary Table S4 and Figure 4. Only associations at *p*<1×10^-15^ are annotated in the image for ease of viewing. Note that a single-SNP Bonferroni correction results in a significance level of 3.3×10^-5^.

**Figure 3:**
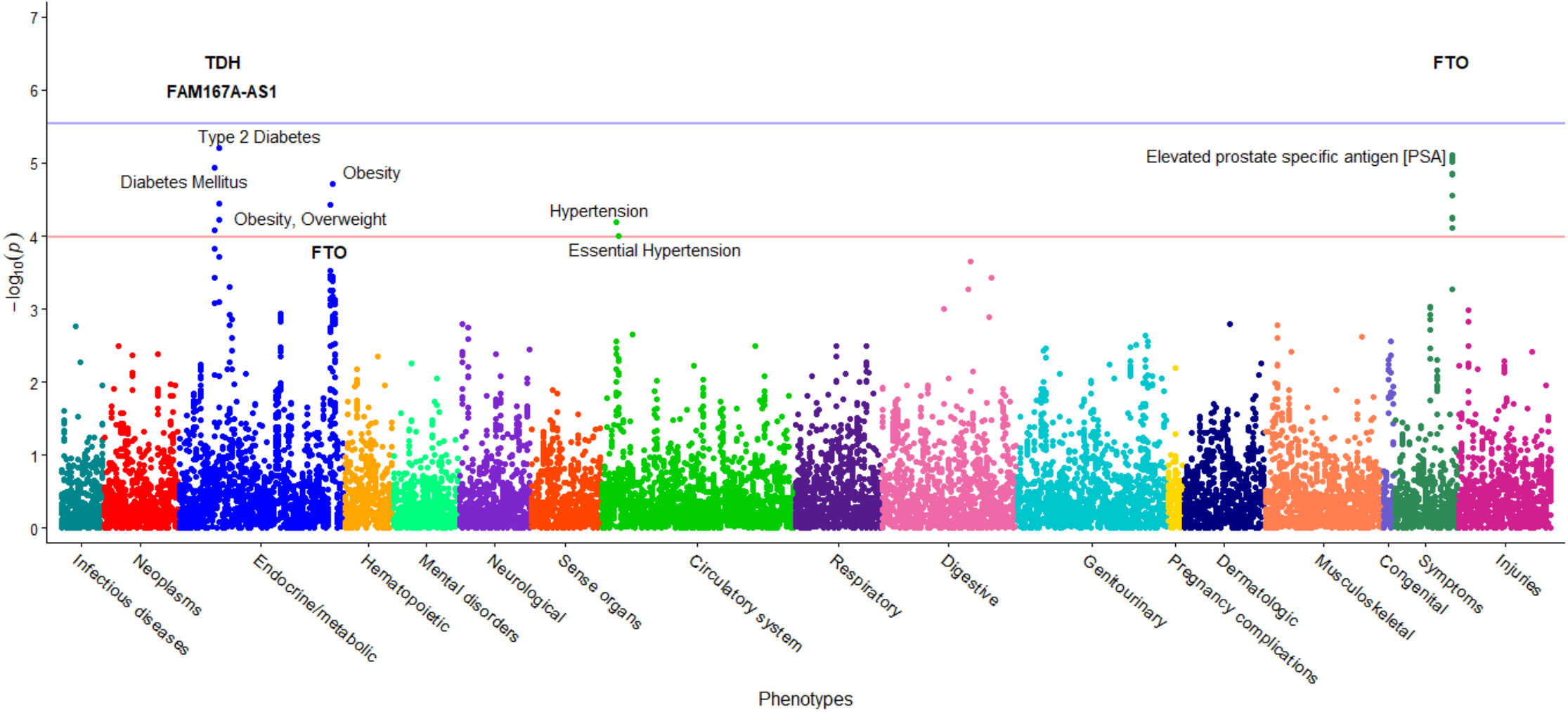
PheWAS results between BMI-significant SNPs and EHR Phenotypes. This figure shows the results of individual logistic regressions between incidence of 633 phenotype groups (phecodes) and the genotypes of 27 SNPs found to have statistically significant associations with BMI in a cohort with DM2 patients. Each point represents the *p*-value of one SNP and one of 633 phecodes with at least 20 cases assigned to it. The horizontal red line represents the significance level *p*=1.02×10^-4^, and the blue line represents the Bonferroni correction of *p*=3×10^-6^.

**Figure 4:**
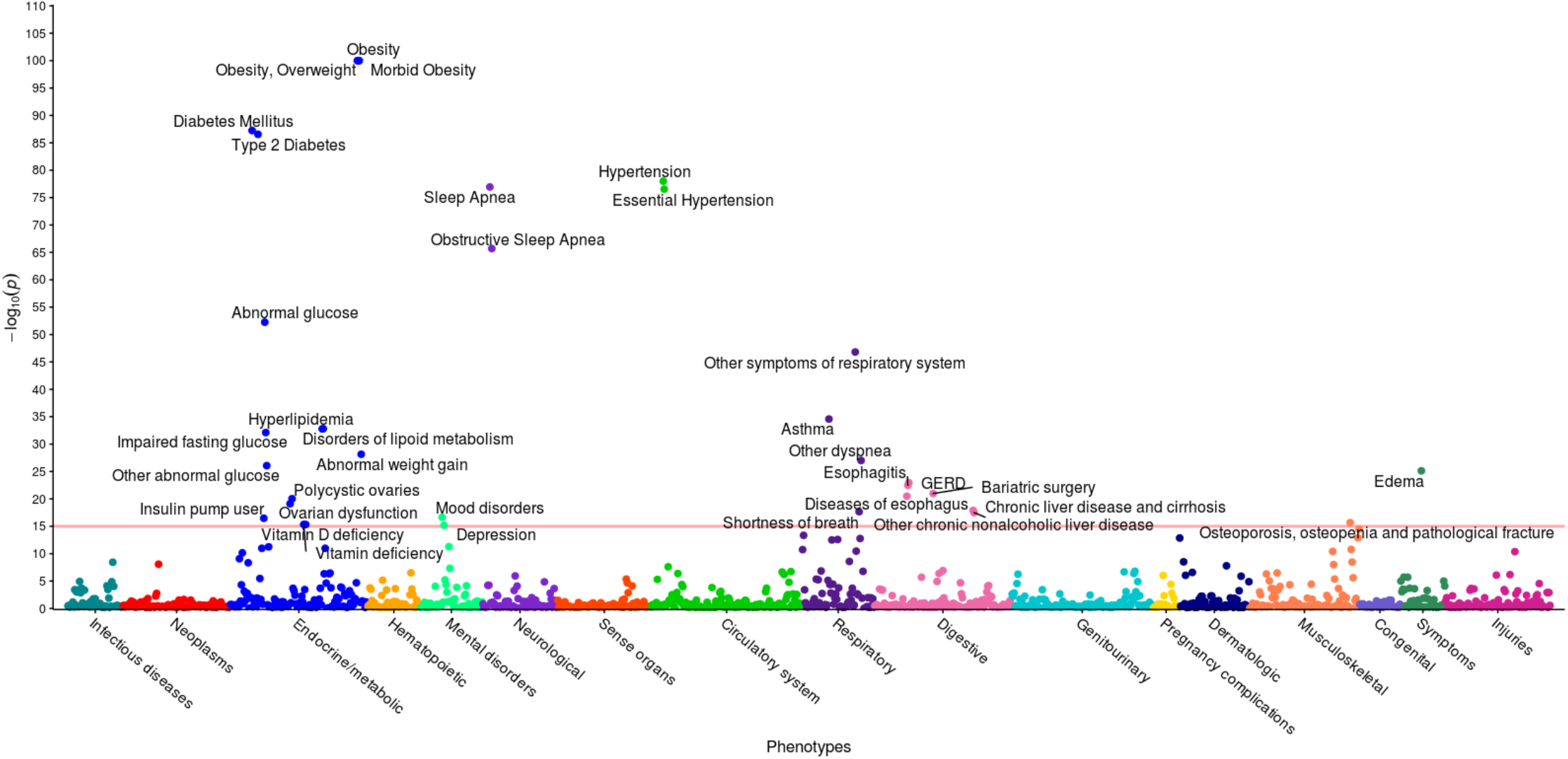
PheWAS results between BMI and EHR Phenotypes. This figure illustrates the results of individual linear regression between incidence of phenotype groups (phecodes) and the continuous BMI metric of all 6,645 individuals. Each of the 301 points represents the *p*-value of the association between one of 1,523 phecodes with at least 20 cases assigned to it, and BMI. Statistical significance was assessed by using the False Discovery Rate of 0.1, corresponding to a raw p-value of 1.96×10^-2^. Only associations with *p* < 1×10^-15^ are annotated for ease of viewing, represented by a horizontal line at 15 on the y-axis.

**Table 5.** Statistically Significant BMI with DM2 PheWAS SNPs.

| Phenotype | Description | SNP | Gene | $\beta$ | (SE) | Odds Ratio | PheWAS *p*-value | N | Cases | Controls |
| --- | --- | --- | --- | --- | --- | --- | --- | --- | --- | --- |
| 250.2 | Type 2 Diabetes | rs2246606 | TDH | -0.331 | 0.073 | 0.718 | 6.32x10<sup>-6</sup> | 5566 | 459 | 5107 |
| 796 | Elevated prostate specific antigen [PSA] | rs9937053 | FTO | 0.882 | 0.197 | 2.416 | 7.76x10<sup>-6</sup> | 6293 | 71 | 6222 |
| 796 | Elevated prostate specific antigen [PSA] | rs9930333 | FTO | 0.881 | 0.197 | 2.413 | 7.88x10<sup>-6</sup> | 6297 | 71 | 6226 |
| 796 | Elevated prostate specific antigen [PSA] | rs9940128 | FTO | 0.875 | 0.197 | 2.398 | 8.66x10<sup>-6</sup> | 6297 | 71 | 6226 |
| 796 | Elevated prostate specific antigen [PSA] | rs1121980 | FTO | 0.872 | 0.197 | 2.391 | 9.60x10<sup>-6</sup> | 6298 | 71 | 6227 |
| 250 | Diabetes Mellitus | rs2246606 | TDH | -0.318 | 0.072 | 0.728 | 1.14x10<sup>-5</sup> | 5577 | 470 | 5107 |
| 796 | Elevated prostate specific antigen [PSA] | rs1421085 | FTO | 0.855 | 0.197 | 2.351 | 1.41x10<sup>-5</sup> | 6298 | 71 | 6227 |
| 796 | Elevated prostate specific antigen [PSA] | rs1558902 | FTO | 0.854 | 0.197 | 2.350 | 1.43x10<sup>-5</sup> | 6298 | 71 | 6227 |
| 278.1 | Obesity | rs2948300 | N/A | 0.199 | 0.047 | 1.221 | 1.90x10<sup>-5</sup> | 5805 | 1172 | 4633 |
| 796 | Elevated prostate specific antigen [PSA] | rs17817449 | FTO | 0.823 | 0.196 | 2.277 | 2.75x10<sup>-5</sup> | 6298 | 71 | 6227 |
| 796 | Elevated prostate specific antigen [PSA] | rs8043757 | FTO | 0.822 | 0.196 | 2.275 | 2.82x10<sup>-5</sup> | 6298 | 71 | 6227 |
| 250.2 | Type 2 Diabetes | rs2572386 | FAM167A | -0.303 | 0.073 | 0.739 | 3.57x10<sup>-5</sup> | 5564 | 459 | 5105 |
| 278 | Obesity, Overweight | rs2948300 | N/A | 0.182 | 0.044 | 1.200 | 3.71x10<sup>-5</sup> | 5980 | 1347 | 4633 |
| 796 | Elevated prostate specific antigen [PSA] | rs8050136 | FTO | 0.788 | 0.195 | 2.198 | 5.55x10<sup>-5</sup> | 6298 | 71 | 6227 |
| 796 | Elevated prostate specific antigen [PSA] | rs9939609 | FTO | 0.787 | 0.196 | 2.197 | 5.71x10<sup>-5</sup> | 6298 | 71 | 6227 |
| 250.2 | Type 2 Diabetes | rs1435277 | N/A | -0.292 | 0.073 | 0.747 | 5.94x10<sup>-5</sup> | 5554 | 458 | 5096 |
| 401 | Hypertension | rs12412241 | N/A | -0.221 | 0.055 | 0.802 | $6.49 \times 10^{-5}$ | 6057 | 1385 | 4672 |
| 796 | Elevated prostate<br>specific antigen [PSA] | rs3751812 | FTO | 0.772 | 0.195 | 2.163 | $7.87 \times 10^{-5}$ | 6297 | 71 | 6226 |
| 250 | Diabetes Mellitus | rs2572386 | FAM167 | -0.284 | 0.072 | 0.753 | $8.44 \times 10^{-5}$ | 5575 | 470 | 5105 |
| 401.1 | Essential<br>Hypertension | rs12412241 | N/A | -0.216 | 0.056 | 0.806 | $1.02 \times 10^{-4}$ | 6037 | 1365 | 4672 |

We performed the same two PheWASs on the case-control GWAS. The first PheWAS identified possible associations between 34 SNPs and 372 phenotype groups with at least 20 individuals (Figure 5). The significance level corresponding to an FDR of 0.1 was *p*=3.85×10^-4^, resulting in 50 significant associations. The Bonferroni correction *p*-value is 4×10^-6^ and shown in blue (see Materials and Methods). The strongest associations occurred between the *NEGR1* and *KLRB1* gene and obesity and morbid obesity and the *NEGR1* gene and abnormal glucose. The *NEGR1* gene, along with *TDH* and *FAM167A-AS1*, also associated with impaired fasting glucose and diabetes, respectively, at a slightly lower, yet still significant *p*-values. The locus rs2948300 on chromosome 8 associated with essential hypertension. The SNP rs1620977 on *NEGR1* is linked with incidence of bronchitis. Additionally, several strong associations of irritable bowel syndrome (IBS) and digestive disorders with loci in *CABP5* are shown. Results are pictured in Figure 5 and included in Table 6.

**Figure 5:**
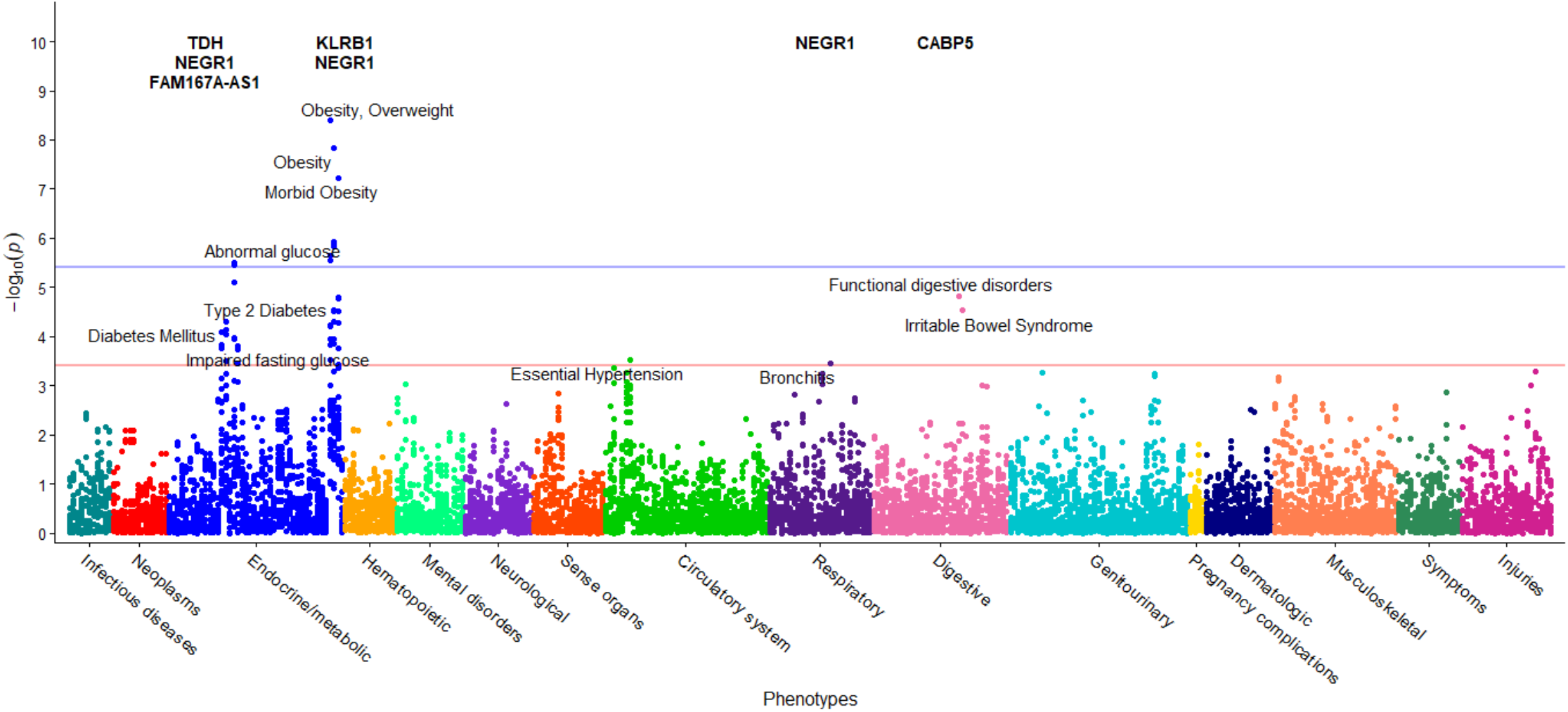
PheWAS results between obesity-significant SNPs and EHR Phenotypes. This figure presents results of individual logistic regressions between incidence of 372 phenotype groups (phecodes) and the genotypes of 34 SNPs found to be associated with extreme obesity. Each point represents the *p*-value of one SNP and one of 372 phecodes with at least 20 cases assigned to it. The horizontal red line represents the significance level *p*=3.85×10^-4^, and the blue line represents the Bonferroni correction of *p*=4×10^-6^.

**Table 6.** Statistically Significant Obesity PheWAS SNPs.

| Phenotype | Description | SNP | Gene | $\beta$ | (SE) | Odds Ratio | PheWAS *p*-value | N | Cases | Controls |
| --- | --- | --- | --- | --- | --- | --- | --- | --- | --- | --- |
| 278 | Overweight, Obesity | rs1620977 | NEGR1 | 0.409 | 0.070 | 1.506 | $4.09 \times 10^{-9}$ | 2819 | 640 | 2179 |
| 278.1 | Obesity | rs1620977 | NEGR1 | 0.405 | 0.071 | 1.499 | $1.45 \times 10^{-8}$ | 2779 | 600 | 2179 |
| 278.11 | Morbid Obesity | rs1620977 | NEGR1 | 0.432 | 0.080 | 1.541 | $5.96 \times 10^{-8}$ | 2624 | 445 | 2179 |
| 278.1 | Obesity | rs1776012 | NEGR1 | -0.326 | 0.067 | 0.722 | $1.18 \times 10^{-6}$ | 2762 | 594 | 2168 |
| 278.1 | Obesity | rs1870676 | NEGR1 | -0.324 | 0.067 | 0.723 | $1.40 \times 10^{-6}$ | 2757 | 593 | 2164 |
| 278.1 | Obesity | rs9424977 | NEGR1 | -0.321 | 0.067 | 0.725 | $1.46 \times 10^{-6}$ | 2776 | 599 | 2177 |
| 278 | Overweight, Obesity | rs1776012 | NEGR1 | -0.308 | 0.065 | 0.735 | $2.23 \times 10^{-6}$ | 2802 | 634 | 2168 |
| 278 | Overweight, Obesity | rs1870676 | NEGR1 | -0.305 | 0.065 | 0.737 | $2.84 \times 10^{-6}$ | 2797 | 633 | 2164 |
| 278 | Overweight, Obesity | rs9424977 | NEGR1 | -0.303 | 0.065 | 0.739 | $2.87 \times 10^{-6}$ | 2816 | 639 | 2177 |
| 250.4 | Abnormal glucose | rs1776012 | NEGR1 | -0.444 | 0.095 | 0.642 | $3.20 \times 10^{-6}$ | 2664 | 272 | 2392 |
| 250.4 | Abnormal glucose | rs1870676 | NEGR1 | -0.444 | 0.096 | 0.642 | $3.58 \times 10^{-6}$ | 2655 | 270 | 2385 |
| 250.4 | Abnormal glucose | rs9424977 | NEGR1 | -0.424 | 0.095 | 0.655 | $7.81 \times 10^{-6}$ | 2674 | 273 | 2401 |
| 564 | Functional digestive disorders | rs750456 | CABP5 | 0.989 | 0.229 | 2.689 | $1.55 \times 10^{-5}$ | 2518 | 40 | 2478 |
| 278.11 | Morbid Obesity | rs2241005 | KLRB1 | -0.321 | 0.075 | 0.725 | $1.64 \times 10^{-5}$ | 2625 | 445 | 2180 |
| 278.11 | Morbid Obesity | rs2058426 | N/A | -0.322 | 0.075 | 0.725 | $1.66 \times 10^{-5}$ | 2621 | 445 | 2176 |
| 278.1 | Obesity | rs2568958 | N/A | -0.294 | 0.070 | 0.745 | $2.89 \times 10^{-5}$ | 2781 | 601 | 2180 |
| 278.1 | Obesity | rs2815752 | N/A | -0.294 | 0.070 | 0.745 | $2.89 \times 10^{-5}$ | 2781 | 601 | 2180 |
| 564.1 | Irritable Bowel Syndrome | rs750456 | CABP5 | 0.992 | 0.238 | 2.697 | $2.98 \times 10^{-5}$ | 2515 | 37 | 2478 |
| 278.11 | Morbid Obesity | rs1870676 | NEGR1 | -0.315 | 0.075 | 0.730 | $3.06 \times 10^{-5}$ | 2605 | 441 | 2164 |
| 278.1 | Obesity | rs3101336 | N/A | -0.293 | 0.070 | 0.746 | $3.08 \times 10^{-5}$ | 2781 | 601 | 2180 |
| 278.11 | Morbid Obesity | rs1776012 | NEGR1 | -0.315 | 0.076 | 0.730 | $3.14 \times 10^{-5}$ | 2606 | 438 | 2168 |
| 250.2 | Type 2 Diabetes | rs2246606 | TDH | -0.468 | 0.115 | 0.626 | $4.98 \times 10^{-5}$ | 2591 | 187 | 2404 |
| 278.1 | Obesity | rs2953802 | N/A | 0.266 | 0.066 | 1.305 | $5.13 \times 10^{-5}$ | 2781 | 601 | 2180 |
| 278.11 | Morbid Obesity | rs9424977 | NEGR1 | -0.304 | 0.075 | 0.738 | $5.21 \times 10^{-5}$ | 2620 | 443 | 2177 |
| 278 | Overweight, Obesity | rs2568958 | N/A | -0.274 | 0.068 | 0.760 | $5.88 \times 10^{-5}$ | 2821 | 641 | 2180 |
| 278 | Overweight, Obesity | rs2815752 | N/A | -0.274 | 0.068 | 0.760 | $5.88 \times 10^{-5}$ | 2821 | 641 | 2180 |
| 278 | Overweight, Obesity | rs3101336 | N/A | -0.273 | 0.068 | 0.761 | $6.27 \times 10^{-5}$ | 2821 | 641 | 2180 |
| 250.2 | Type 2 Diabetes | rs2736280 | TDH | -0.452 | 0.114 | 0.636 | $7.48 \times 10^{-5}$ | 2588 | 187 | 2401 |
| 250 | Diabetes Mellitus | rs2246606 | TDH | -0.447 | 0.114 | 0.639 | $8.11 \times 10^{-5}$ | 2596 | 192 | 2404 |
| 250.2 | Type 2 Diabetes | rs2953802 | N/A | 0.431 | 0.110 | 1.539 | $8.90 \times 10^{-5}$ | 2591 | 187 | 2404 |
| 250.4 | Abnormal glucose | rs2568958 | N/A | -0.389 | 0.100 | 0.678 | $1.08 \times 10^{-4}$ | 2678 | 274 | 2404 |
| 250.4 | Abnormal glucose | rs2815752 | N/A | -0.389 | 0.100 | 0.678 | $1.08 \times 10^{-4}$ | 2678 | 274 | 2404 |
| 278 | Overweight, Obesity | rs2241005 | KLRB1 | -0.247 | 0.064 | 0.781 | $1.11 \times 10^{-4}$ | 2821 | 641 | 2180 |
| 250.4 | Abnormal glucose | rs3101336 | N/A | -0.388 | 0.101 | 0.678 | $1.12 \times 10^{-4}$ | 2678 | 274 | 2404 |
| 278.1 | Obesity | rs2241005 | KLRB1 | -0.253 | 0.066 | 0.776 | $1.14 \times 10^{-4}$ | 2781 | 601 | 2180 |
| 278.1 | Obesity | rs2058426 | N/A | -0.250 | 0.066 | 0.779 | $1.43 \times 10^{-4}$ | 2777 | 601 | 2176 |
| 278 | Overweight, Obesity | rs2058426 | N/A | -0.243 | 0.064 | 0.784 | $1.46 \times 10^{-4}$ | 2817 | 641 | 2176 |
| 250 | Diabetes Mellitus | rs2736280 | TDH | -0.426 | 0.112 | 0.653 | $1.48 \times 10^{-4}$ | 2593 | 192 | 2401 |
| 250.41 | Impaired fasting glucose | rs1870676 | NEGR1 | -0.471 | 0.125 | 0.625 | $1.59 \times 10^{-4}$ | 2535 | 150 | 2385 |
| 278.11 | Morbid Obesity | rs2953802 | N/A | 0.278 | 0.074 | 1.321 | $1.76 \times 10^{-4}$ | 2625 | 445 | 2180 |
| 250 | Diabetes Mellitus | rs2953802 | N/A | 0.407 | 0.109 | 1.502 | $1.77 \times 10^{-4}$ | 2596 | 192 | 2404 |
| 250.41 | Impaired fasting glucose | rs1776012 | NEGR1 | -0.464 | 0.124 | 0.629 | $1.80 \times 10^{-4}$ | 2543 | 151 | 2392 |
| 401.1 | Essential Hypertension | rs2948300 | N/A | 0.282 | 0.078 | 1.326 | $3.02 \times 10^{-4}$ | 2737 | 521 | 2216 |
| 278 | Overweight, Obesity | rs2953802 | N/A | 0.231 | 0.064 | 1.260 | $3.06 \times 10^{-4}$ | 2821 | 641 | 2180 |
| 250.2 | Type 2 Diabetes | rs2572386 | FAM167A | -0.415 | 0.115 | 0.660 | $3.15 \times 10^{-4}$ | 2590 | 187 | 2403 |
| 497 | Bronchitis | rs1620977 | NEGR1 | 0.673 | 0.188 | 1.960 | $3.47 \times 10^{-4}$ | 2118 | 60 | 2058 |
| 250.41 | Impaired fasting glucose | rs9424977 | NEGR1 | -0.441 | 0.124 | 0.644 | $3.60 \times 10^{-4}$ | 2552 | 151 | 2401 |
| 278.11 | Morbid Obesity | rs2568958 | N/A | -0.283 | 0.079 | 0.753 | $3.66 \times 10^{-4}$ | 2625 | 445 | 2180 |
| 278.11 | Morbid Obesity | rs2815752 | N/A | -0.283 | 0.079 | 0.753 | $3.66 \times 10^{-4}$ | 2625 | 445 | 2180 |
| 278.11 | Morbid Obesity | rs3101336 | N/A | -0.282 | 0.080 | 0.754 | $3.85 \times 10^{-4}$ | 2625 | 445 | 2180 |

The second PheWAS in this case-control study identified possible links between 1,362 EHR phenotype groups with at least 20 individuals and the incidence of extreme obesity (Figure 6). The significant threshold of *p*=1.4×10^-2^ enabled an FDR of 0.1. At this significance level, 191 significant associations were identified, including obesity (*p*<1×10^-134^), hypertension *(p*<1×10^-65^), sleep apnea (*p* < 1×10^-43^), abnormal glucose (*p*<1×10^-40^), hyperlipidemia, asthma, and GERD. The high level of association with obesity validates our methods. These associations are shown in Figure 6, with phenotypes annotated above a significance level of 1×10^-15^ for ease of viewing. A line is drawn at the significance level 1×10^-15^ as guidance. Results are included in Supplementary Table S5. Note that a single-SNP Bonferroni correction results in a significance level of 3.7×10^-5^.

**Figure 6:**
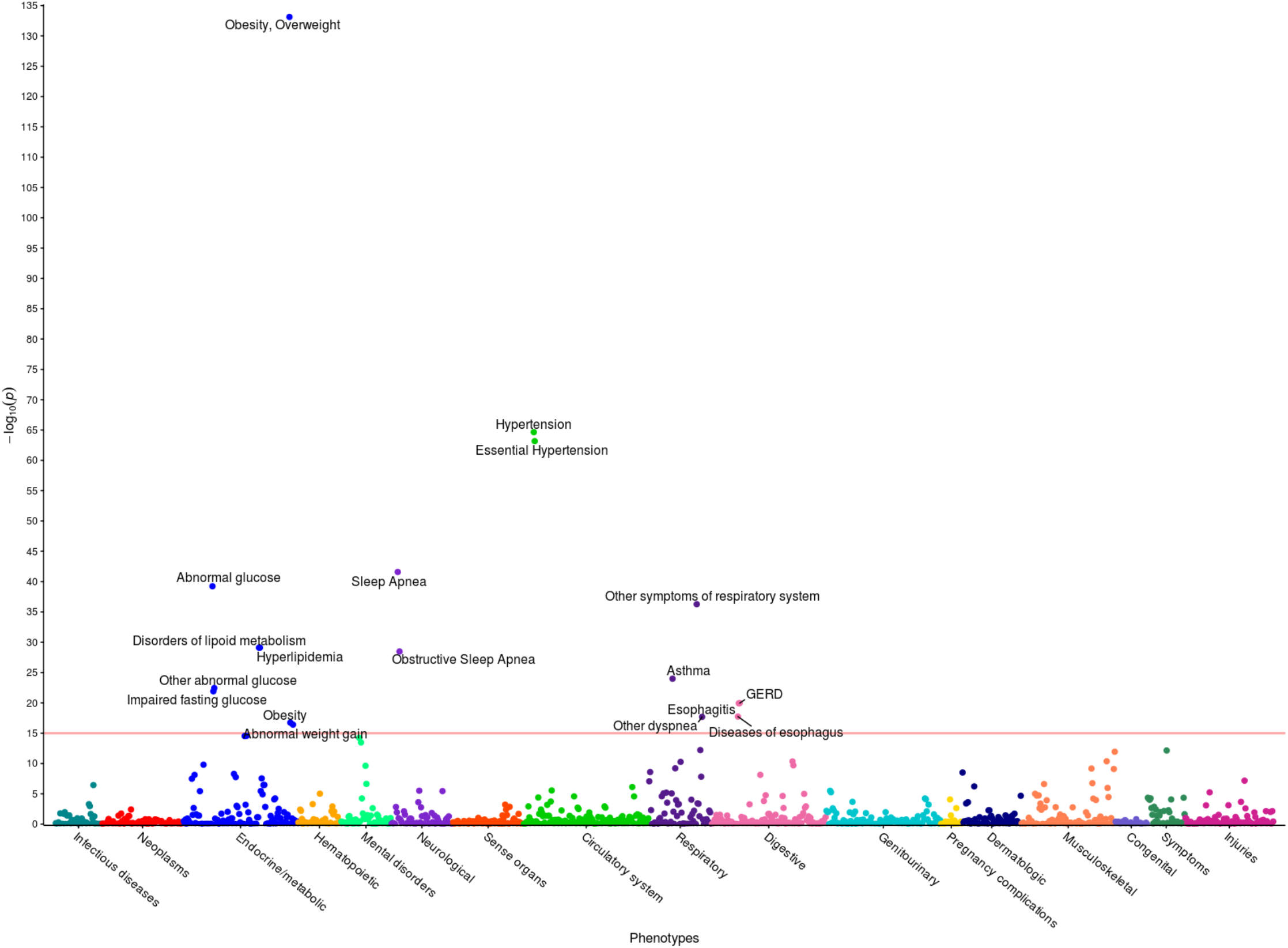
PheWAS results between extreme obesity and EHR Phenotypes. This figure illustrates the results of individual linear regression between incidence of phenotype groups (phecodes) and the incidence of extreme obesity in 2,996 individuals. Each of the 191 points represents the *p*-value of the association between one of 1,362 phecodes with at least 20 cases assigned to it, and extreme obesity. Statistical significance was assessed by using the False Discovery Rate of 0.1, corresponding to a raw p-value of 1.4×10^-2^. Only associations with *p* < 1×10^-15^ are annotated for ease of viewing, represented by a horizontal line at 15 on the y-axis.

## Discussion

### GWAS of Healthy Nevada BMI and Obesity

Here we present three GWASs on participants in the Healthy Nevada Project. The first investigates associations with BMI on subjects that do not have DM2 diagnoses. The second identifies associations with BMI in which participants with DM2 are included as a comorbidity. The third GWAS is a case-control study of extreme obesity that complements outcomes of the first two quantitative trait studies. Each GWAS is followed with two independent PheWASs to examine pleiotropy and additional phenotypic associations with quantitative BMI levels and incidence of obesity.

The first GWAS tested the association between genotype and BMI without DM2, to remove DM2 effects on BMI. As expected, the majority of the resulting associations were found in SNPs that lie within the *FTO* gene. This gene has been associated to BMI and obesity in several studies and is a major focal point in obesity-related research (Scuteri *et al*. 2007; Frayling *et al*. 2007; Dina *et al*. 2007; Zeggini *et al*. 2007; Hinney *et al*. 2007; Hunt *et al*. 2008; Price *et al*. 2008; Grant *et al*. 2008; Hotta *et al*. 2008; Loos *et al*. 2008; Tan *et al*. 2008; Villalobos-Comparán *et al*. 2008; Thorleifsson *et al*. 2008; Willer *et al*. 2009; Meyre *et al*. 2009; Wing *et al*. 2009; Shimaoka *et al*. 2010; Fawcett and Barroso 2010; Wang *et al*. 2011; Prakash *et al*. 2011; Okada *et al*. 2012; Berndt *et al*. 2013; Wheeler *et al*. 2013; Graff *et al*. 2013; Olza *et al*. 2013; Qureshi *et al*. 2017; Gonzalez-Herrera *et al*. 2018). Moreover, the two strongest associations in the GWAS were from SNPs in *FTO* (rs1558902/ rs1421085, *p* = 1.67×10^-7^/1.75×10^-7^), highlighting the overall importance of this gene in relation to obesity. Frayling suggests that the association of *FTO* SNPs with DM2 is mediated through BMI (Frayling *et al*. 2007).

The exact mechanism by which the *FTO* gene affects BMI is not understood; however, it has been discovered that the gene product of *FTO* mediates oxidative demethylation of several different RNA species, such as mRNA, snRNA and tRNA (Jia *et al*. 2011; Wei *et al*. 2018). This indicates that protein produced from *FTO* likely operates as a RNA regulatory molecule, which can affect both gene expression as well as translation initiation and elongation (Wei *et al*. 2018).

Two SNPs within *TDH* were found to be strongly associated to BMI. This gene codes for a nonfunctional L-threonine dehydrogenase, lacking most of the C-terminus found in other species, and is thus characterized as a putative pseudogene. Previous research has identified this gene as a possible susceptibility gene for obesity (Yanagiya *et al*. 2007); however, relatively little is known about any functional consequences of SNPs within this pseudogene. We also observed a strongly associated locus in *NEGR1*, one of the first genes shown to have variants associated to BMI (Thorleifsson *et al*. 2008; Willer *et al*. 2009; Speliotes *et al*. 2010; Boender *et al*. 2014). This gene codes for a cell adhesion molecule, although its function in relation to BMI is still unknown. Previous research in mice determined that deletions of *NEGR1* cause a decrease in weight and a change in the regulation of energy balance, implying that *NEGR1* most likely functions to control the regulation of energy balance (Lee *et al*. 2012; Boender *et al*. 2014).

We hypothesized that including DM2 participants (and thus DM2 as a covariate in the genetic model) would produce a more parsimonious fit, as many studies show a relationship between diabetes and BMI. We discovered that diabetes was indeed an important predictor of BMI for all 500,000 regressions performed (*p* < 2×10^-16^ for every SNP), regardless of age, gender or genotype. Furthermore, adding DM2 as a predictor in the additive model increases the significance of associations between SNPs in the *TDH* gene and BMI; five out the top eight most significant associations fall within this gene (Table 2). It is clear that incidence of DM2 in our cohort affects the genetic association of BMI. Specifically, when DM2 patients are excluded in our cohort, there are two associations within *TDH*. When DM2 participants are included, we observe five associations with the *TDH* gene. This indicates, along with the BMI PheWAS results, that not only does *TDH* influence BMI measurements, it also has an association with DM2.

It is rare for SNPs to be effectors of two separate diseases, even those as intertwined as BMI and DM2 (Grarup *et al*. 2014). A possible explanation why *TDH* has not been previously associated with DM2 is due to a lack of statistical power to observe the small risk increases *TDH* may impose on DM2 (Grarup *et al*. 2014). Nonetheless, the increased rate of DM2 diagnosis worldwide makes this an interesting candidate gene. How the SNPs in the *TDH* pseudogene may influence either BMI or DM2 is unknown, as evidence supporting the association between *TDH* and BMI/DM2 is scant; however, previous research has discovered that not all pseudogenes are “junk” DNA. Some of these genes can be actively transcribed to produce short interfering RNAs (siRNAs), which can regulate gene expression (Pink *et al*. 2011). In certain cases, they can even competitively bind micro-RNAs (miRNAs), which can attenuate repression of cellular mRNA (An *et al*. 2016). Additionally, the expression of pseudogene transcripts tends to be tissue-specific. Given that the greatest expression of *TDH* transcripts is found in the pancreas (Fagerberg *et al*. 2014), one might speculate that *TDH* affects the production of insulin and/or digestive enzymes. If true, this may account for our observation where *TDH* influences BMI measurements, and is associated with DM2. Given the strong associations between *TDH* SNPs and DM2, as well as potential regulatory functions of pseudogenes, we believe it is essential that future studies focus on determining the function of the *TDH* pseudogene in a tissue-specific context. Although studies using genetically modified mice with a *TDH* polymorphism and proteomics analysis of their pancreatic tissue would be straight-forward, to the best of our knowledge, no such studies have been reported. We see this as a possible future direction.

Adding DM2 as a covariate into the statistical model also identified two additional genes that may influence BMI: *RTN4RL1* and *CABP5*. *RTN4RL1* is a gene that codes for a cell surface receptor and was previously found to be upregulated approximately 2-fold when exposed to bone morophogenetic protein 4 (BMP4), a protein that is increased in diabetic animals and may reduce insulin secretion (Christensen *et al*. 2015). This implies that the effects of *RTN4RL1* on BMI may be secondary to its main effect on diabetes. Moreover, this gene has also been listed as a potential candidate gene for DM2 in previous GWAS (Thomsen *et al*. 2016). The gene *CABP5*, which codes for a calcium binding protein that has role in calcium mediated cellular signal transduction (Haeseleer *et al*. 2002) may have a more direct effect on BMI. It was previously discovered as part of a group of several genes that were upregulated in obese individuals, although its exact function relative to obesity is still unknown (Nakajima *et al*. 2016).

A case-control association study examining the effects between genetics and the risk of extreme obesity (BMI ≥ 35 kg/m^2^) was the final GWAS we conducted. It has been determined by the World Health Organization (WHO) that more than 1.9 billion adults are overweight and over 650 million are obese. Moreover, obesity is associated with several other chronic diseases, such as cardiovascular disease, DM2 and cancer, all of which could lead to premature death (Kopelman 2007). Overall, our obesity results consist of many of the same SNPs and genes found to be associated with BMI. However, the obesity results did demonstrate an increase in the genetic associations at the significance level of *p*<1×10^-5^ in and very close to the *NEGR1* gene compared to previous BMI associations. Previous studies using genetically modified mice with *NEGR1* deficiency or *NEGR1*-loss-of-function support a role for *NEGR1* in the control of body weight; however, the mechanism of its involvement is not clear (Lee *et al*. 2012). Contradictory with anticipated results, these mutant mice display a small but steady reduction of body mass. Notwithstanding, these studies do suggest that loss of *NEGR1* function in the mouse models has a negative effect on body mass as well as lean mass, supporting the possibility that *NEGR1* may contribute to a change body mass. It is also important to note that animal models are a representation of human physiology but not necessarily a precise depiction.

Our extreme obesity vs. non-obese study also identified two new genes, *PFKFB3* and *KLRB1*, that are not yet found to be significantly associated to BMI. The odds ratios associated with all SNPs found in these genes were less than one, indicating that they decreased the odds of extreme obesity risk. These results are potentially supported by work conducted by Huo et al., who reported that mice transgenically modified to selectively overexpress *PFKFB3* in adipocytes show increases fat deposition in their adipose tissue (Huo *et al*. 2012). In contrast, an earlier study reported that transgenic mice with reduced *PFKFB3* expression show exacerbated diet-induced insulin resistance (Huo *et al*. 2010). Moreover, a recent study of hypertrophic white adipose tissue morphology, Kerr and coworkers showed that inhibition of *PFKFB3* mRNA impairs basal and insulin stimulated lipogenesis and furtherer proposed that gene knockdown may of *PFKFB3* inhibit adipocyte lipid storage (Kerr *et al*. 2019). These studies support a hypothesis whereby polymorphisms that lead to a decrease in *PFKFB3* may be protective from the development of obesity; however, tissue-specific transcriptional studies in humans would be required to fully support this assertion.

We also observed polymorphisms in *KLRB1* to associate with a decrease of odds in extreme obesity risk. *KLRB1* expression produces a type II transmembrane glycoprotein also known as CD161; a member of the C-type lectin superfamily. CD161 is expressed on the surface of most natural killer (NK) cells and natural killer T (NKT) but also on subsets of peripheral T cells and CD3^+^ thymocytes. While the biological function of CD161 is not firmly established, it was suggested that it serves either as a stimulatory receptor or to inhibit NK cell-mediated cytotoxicity and cytokine production (Lanier *et al*. 1994). Indeed, NK cells were shown to be upregulated in the fat of obese twins (Pietiläinen *et al*. 2008); moreover, BMI and *KLRB1* expression may be correlated in that *KLRB1 transcription has been reported to increase* as BMI increases (Rai *et al*. 2014). Additionally, CD161^bright^ CD8^+^ mucosal associated invariant T (MAIT) cells play a central role in maintaining mucosal immunity and therefore, changes in CD161 expression on these cells may lead to alterations in mucosal immunity and gut microbiota homeostasis. These changes may in turn manifest as alterations of dietary metabolism. It is noteworthy that increases in MAIT cells are associated with Juvenile Type 1 Diabetes and polymorphisms in *KLRB1* have been associated with ischemic heart disease (Makeeva *et al*. 2015), and differential transcription of *KLRB1* has been reported in DM2 and coronary artery disease (Gong *et al*. 2017). Furthermore, another gene, *GRIN2A*, is a gene that is part of the family of genes {*GRIN1*, *GRIN2A*, *GRIN2B*, *GRIN2C*, *GRIN2D*, *GRIN3A*, and *GRIN3B*}, which encode proteins that form a receptor in charge of sending chemical messages between neurons in the brain. The gene *GRIN2B* was found to be associated to obesity in adult women defined as metabolically healthy in Schlauch et. al (*p*=1.7×10^-5^). (Schlauch *et al*. 2019).

Associations between *FTO* and obesity were just under genome-wide significance levels. This is a possible indication that *FTO* polymorphisms cause small changes in BMI, rather than the wide range differences observed between extreme obese cases and controls. Nonetheless, previous research has demonstrated that a combination of several *FTO* mutations will increase the likelihood of a participant being classified as obese (Li *et al*. 2009). Speakman et al. stated that Frayling showed the *FTO* was significantly associated with diabetes only through its association with BMI (Speakman *et al*. 2018).

The comprehensive series of GWASs presented here validates associations of obesity and BMI found in previous studies, such as the *FTO* and *NEGR1* loci (Willer *et al*. 2009; Speliotes *et al*. 2010; Okada *et al*. 2012; Locke *et al*. 2015). Many larger studies identify associative loci in *MC4R* (Willer *et al*. 2009; Speliotes *et al*. 2010; Okada *et al*. 2012). While our studies did not detect SNPs in *MC4R* with genome-wide significance, they did identify associations at *p*=1×10^-4^ and *p*=1.7×10^-4^ of SNPs rs17782313 and rs571312 (Willer *et al*. 2009; Speliotes *et al*. 2010). A number of obesity case-control studies have found variations in the *MC4R* gene (Xi *et al*. 2012; Evans *et al*. 2014). Our case-control study does reveal that rs17782313 in *MC4R* associates with obesity at *p*=3×10^-4^. Our cohort is a controlled, regional population. The next two stages of the Healthy Nevada Project will add between 40,000 (2019) and 150,000 (late 2020) more Nevadans to the current cohort. With these much larger cohort sizes, it is our hope that a stronger associate link with *MC4R* will be identified.

### PheWAS of Healthy Nevada BMI and Obesity

To the best of our knowledge, this is the first dual-PheWAS targeted at BMI and obesity. Cronin et al. present a comprehensive PheWAS targeted at *FTO* variants, which also show strong associations with overweight and obesity phenotypes, hypertension and hyperlipidemia (Cronin *et al*. 2014). Milliard et al. perform a large PheWAS study to examine phenotypic associations with BMI that focus on the nervousness phenotypes: the study identified known associations such as diabetes and hypertension (Millard *et al*. 2019).

The PheWAS performed on the SNP associations in this study’s BMI cohort identified strong associations of elevated PSA levels with variants in *FTO*, and indicates that the number of minor alleles of these variants is predictive of elevated PSA. This finding is in contradiction to the reports (Bañez *et al*. 2007; Oh *et al*. 2013; Zhang *et al*. 2016; Bonn *et al*. 2016) indicating an inverse relationship between PSA levels and BMI. However, serum levels of PSA may be elevated due to reasons other than prostatic malignancy. Benign prostatic hyperplasia (BPH), prostatitis (Nadler *et al*. 1995), ejaculation (Herschman *et al*. 1997), or manipulation of the prostate gland (Chybowski *et al*. 1992; Crawford *et al*. 1992; Tarhan *et al*. 2005) may cause elevated levels of serum PSA. Our study did not control for such parameters. Our sample includes only 71 individuals with ICD codes indicating high PSA levels, of which a number are morbidly obese. Increased BMI is often associated with increased age and our study population was significantly older than the general median age of the U.S. population. Thus, it is possible that older age contributed to increased likelihood of BPH and, subsequently, elevated serum PSA, negating the reported inverse effect of BMI on PSA levels and possibly exposing a novel association with variants in *FTO*.

Many of the clinical associations observed in the PheWASs of the HNP in relation to various degrees of increased BMI and the presence of obesity are recognizable of the cluster of clinical conditions associated with metabolic syndrome (Alberti *et al*. 2009). Obesity is a risk factor for respiratory conditions such as chronic obstructive pulmonary disease (COPD), asthma, obstructive sleep apnea and obesity hypoventilation syndrome, and may influence the development and presentation of these diseases (Poulain *et al*. 2006). Accumulation of fat tissue impairs ventilatory function in adults (Lazarus *et al*. 1997) and increased BMI is associated with a reduction in forced expiratory volume in one second (FEV1), forced vital capacity (FVC), total lung capacity, functional residual capacity and expiratory reserve volume (Rubinstein *et al*. 1990; Chinn *et al*. 1996; Lazarus *et al*. 1997; Biring *et al*. 1999). Peripheral edema has long been recognized as associated with extreme obesity (Alexander *et al*. 1962). In the U.K. Community Nursing Services study, obesity was found as an independent risk factor for chronic edema (Moffatt *et al*. 2019).

## Data Availability Statement

### EHR Data

EHR data for the Healthy Nevada cohort are subject to HIPAA and other privacy and compliance restrictions. Mean quality-controlled BMI values for the 6,645 component values for each individual are available in Supplementary Table S6.

### GWAS Results

To reduce the possibility of a privacy breach, 23andMe requires that the statistics for only 10,000 SNPs be made publicly available. This is the amount of data considered by 23andMe to be insufficient to enable a reidentification attack. The statistical summary results of the top 10,000 SNPs for the 23andMe data are available here: www.dri.edu/HealthyNVProjectGenetics. All column definitions are listed in Table 7.

**Table 7.** Column Identifiers for GWAS Results.

| Column name | Definition |
| --- | --- |
| CHR | Chromosome |
| SNP | Individual SNP identifier |
| BP | Location of SNP on relative chromosome |
| A1 | Alternative Allele |
| TEST | Selected statistical test – ADD represents the additive effect |
| NMISS | Indicates the number of observations – non-missing genotypes |
| BETA | The effect size for this variant, defined per copy of the A1 allele |
| SE | The standard error of the effect size |
| LE | Lower end of the 95% confidence interval for the effect size |
| UE | Upper end of the 95% confidence interval for the effect size |
| STAT | The value of the test statistic |
| P | The p-value for the association test |

Table describing the column headers for the results file of our genome wide associations. This summary results file only lists the top 10,000 SNPs in order to prevent a re-identification attack.

### PheWAS Results

Summarized counts of each ICD classification (ICD-9 and ICD-10) and phenotype group (phecode) are presented in Supplementary Table S7.

## Funding statement and role of funders

Research support was provided by the Governor’s Office of Economic Development Knowledge Fund. Support for the Healthy Nevada Project and personal genetics was provided by the Renown Health and the Renown Health Foundation. 23andMe provided support of salaries for members of the 23andMe Research Team, who are employees of 23andMe, Inc. None of the funders had any role in the study design, data collection and analysis, decision to publish, or preparation of the manuscript. The specific roles of these authors are articulated in the Author Contributions section.

## Competing interest

Members of the 23andMe Research Team are employees of 23andMe and hold stock or stock options in 23andMe. 23andMe provided support in the form of salaries for members of the 23andMe Research Team. Affiliation with 23andMe does not alter our adherence to Genetics’ policies on sharing data and materials. Please see the Data availability section above.

## Acknowledgements

We thank Michele Henderson, Toni Curreri and all the ambassadors of the Healthy Nevada Project. We thank Renown Health and DRI marketing and all the folks at 23andMe who helped launch the project. Research support was provided by the Governor’s Office of Economic Development Knowledge Fund. Support for the Healthy Nevada Project and personal genetics was provided by the Renown Health Foundation.

Members of the 23andMe Research Team are: Michelle Agee, Adam Auton, Robert K. Bell, Robert Borkowski, Katarzyna Bryc, Sarah L. Elson, Pierre Fontanillas, Nicholas A. Furlotte, David A. Hinds, Karen E. Huber, Aaron Kleinman, Nadia K. Litterman, Matthew H. McIntyre, Joanna L. Mountain, Elizabeth S. Noblin, Carrie A.M. Northover, Steven J. Pitts, J. Fah Sathirapongsasuti, Olga V. Sazonova, Janie F. Shelton, Suyash Shringarpure, Chao Tian, Joyce Y. Tung, Vladimir Vacic, and Catherine H. Wilson.

## Supplemental Figure and Table Legends

**Supplementary Table S1: GWAS results for BMI in a cohort with no DM2-diagnosed participants** This table lists the 20 statistically significant SNPs associated with BMI in our cohort without DM2-diagnosed individuals. General information about the SNP such as chromosome location, GWAS *p*-value, power, genotype, cytoband, and ANOVA are listed.

**Supplementary Table S2: GWAS results for BMI in a cohort with DM2-diagnosed participants** This table lists the 27 statistically significant SNPs associated with the BMI in our cohort that includes all individuals with DM2. General information about the SNP such as chromosome location, GWAS *p*-value, power, genotype, cytoband, and ANOVA are listed.

**Supplementary Table S3: GWAS results for the extreme obesity case-control study** This table lists the 26 statistically significant SNPs associated with extreme obesity. General information about the SNP such as chromosome location, GWAS *p*-value, power, genotype, cytoband, and ANOVA are listed.

**Supplementary Table S4: Significant EHR phenotypic associations with BMI** This is a table of the 301 phenotype groups (phecodes) reaching statistical significance (*p<*1.96×10^-2^) when associated to BMI in our cohort including DM2-diagnosed individuals. Phecodes and their description, effect sizes (β) of the regression, standard error (SE), and *p*-values are included. Each phecode group contains at least 20 cases.

**Supplementary Table S5: Significant EHR phenotypic associations with extreme obesity** This is a table of the 191 phenotype groups (phecodes) reaching statistical significance (*p<*1.4×10^-2^) when associated to extreme obesity in our cohort. Phecodes and their description, effect sizes (β) of the regression, standard error (SE), and *p*-values are included. Each phecode group contains at least 20 cases.

**Supplementary Table S6: Quality-controlled and averaged BMI participant values** This table includes the quality-controlled average BMI value across multiple records for each individual, as well as age at Jan 2019 and gender. Due to the length of this table, it can be found at: www.dri.edu/HealthyNVProjectGenetics

**Supplementary Table S7: Counts of each phecode group** This table presents the mapping between ICD codes (ICD-9 and ICD-10) and phecodes as presented in Carroll and the **R** package PheWAS (Carroll *et al*. 2014) tested in our study, and the number of incidences from the BMI cohort in each phecode group. The column labled Count is derived by the aggreagation of ICD-9 and ICD-10 codes.

**Supplementary Figure S1: Quality-controlled average BMI values of participants** This figure illustrates both the raw and normalized BMI values. Due to certain extreme BMI values, there is a break in the x-axis of the raw BMI values shown in the first panel.

**Supplementary Figure S2: Manhattan plot of GWAS results of BMI with DM2-diagnosed individuals removed** Genome-wide association study results for BMI. This study excludes DM2-diagnosed individuals. The *x*-axis represents the genomic position of 500,508 SNPs. The *y*-axis represents -log_10_-transformed raw *p*-values of each genotypic association. The red horizontal line indicates the significance level 1×10^-5^.

**Supplementary Fig S3: ANOVA results for rs9939609** A box and whisker figure of ANOVA results for one of the strongest associations (rs9939609) with BMI is shown in Supplementary Figure S3. Note that BMI levels increase with the increase of the number of minor alleles, which is typical of variants in *FTO* (Frayling *et al*. 2007).

## Author Contributions

KAS and RWR conducted genetic and clinical data analysis and wrote the manuscript. GE, ADS and KAS contributed to the clinical discussion. VCL contributed to the molecular biology discussion. WJM extracted participants and their clinical health data from the Renown EHR. The 23andMe research team provided participant genotype data and edited the manuscript. JJG obtained funds to conduct this experiment, provided leadership in all aspects of the research, and edited the manuscript. All authors reviewed, edited and approved the final version of the manuscript.

